# Deconstructing intratumoral heterogeneity through multiomic and multiscale analysis of serial sections

**DOI:** 10.1101/2023.06.21.545365

**Authors:** Patrick G. Schupp, Samuel J. Shelton, Daniel J. Brody, Rebecca Eliscu, Brett E. Johnson, Tali Mazor, Kevin W. Kelley, Matthew B. Potts, Michael W. McDermott, Eric J. Huang, Daniel A. Lim, Russell O. Pieper, Mitchel S. Berger, Joseph F. Costello, Joanna J. Phillips, Michael C. Oldham

## Abstract

Tumors may contain billions of cells including distinct malignant clones and nonmalignant cell types. Clarifying the evolutionary histories, prevalence, and defining molecular features of these cells is essential for improving clinical outcomes, since intratumoral heterogeneity provides fuel for acquired resistance to targeted therapies. Here we present a statistically motivated strategy for deconstructing intratumoral heterogeneity through multiomic and multiscale analysis of serial tumor sections (MOMA). By combining deep sampling of IDH-mutant astrocytomas with integrative analysis of single-nucleotide variants, copy-number variants, and gene expression, we reconstruct and validate the phylogenies, spatial distributions, and transcriptional profiles of distinct malignant clones. By genotyping nuclei analyzed by single-nucleus RNA-seq for truncal mutations, we further show that commonly used algorithms for identifying cancer cells from single-cell transcriptomes may be inaccurate. We also demonstrate that correlating gene expression with tumor purity in bulk samples can reveal optimal markers of malignant cells and use this approach to identify a core set of genes that is consistently expressed by astrocytoma truncal clones, including *AKR1C3*, whose expression is associated with poor outcomes in several types of cancer. In summary, MOMA provides a robust and flexible strategy for precisely deconstructing intratumoral heterogeneity and clarifying the core molecular properties of distinct cellular populations in solid tumors.

## Introduction

Tumors are complex ecosystems containing huge numbers of malignant and nonmalignant cells. Malignant cells evolve over time by acquiring mutations through diverse mechanisms that promote genetic^1^ and epigenetic^2^ heterogeneity, which may occur in a neutral fashion^3^ or as a Darwinian response to therapeutic or other environmental pressures^4^. Nonmalignant cells comprise diverse tumor microenvironments (TMEs) that vary within and among tissues and individuals and may be influenced by malignant cells to adopt tumor-suppressive or tumor-supportive behaviors^5,6,7^. The genetic, epigenetic, and microenvironmental diversity of individual tumors is collectively described as intratumoral heterogeneity (ITH)^8^. Clarifying the extent of ITH is an important goal for precision medicine, since most mutations are not shared between malignant clones from different individuals^9–13^ and ITH provides the substrate for acquired resistance to targeted therapies^8,14–16^.

Investigators have mostly studied ITH by applying multiomic assays to a small number of bulk subsamples from the same tumor. Multi-region analyses of renal carcinoma^17^, breast cancer^18^, colorectal cancer^19^, glioblastoma^20^, and others^21^ have identified spatial variation in mutation frequencies and other molecular phenotypes, revealing extensive ITH. However, this experimental design is high-dimensional in omics feature space but low-dimensional in sample space, which can lead to biased inference and inflated false-positive error rates for molecular features^22^. Furthermore, the small number of bulk samples limits conclusions that can be drawn about distinct malignant clones and nonmalignant cell types. Recent efforts using single-cell methods have provided new perspectives on ITH^23–25^, but it remains non-trivial to isolate and sequence DNA and RNA from the same cell at scale. As such, cancer cells are often identified from copy-number variants (CNVs) inferred from single-cell data. However, single-cell data are confounded by technical factors related to tissue dissociation, sampling bias, noise, contamination, and sparsity^26–30^, which muddle the relationships between malignant cell genotypes and molecular phenotypes, particularly for cancers that lack consistent CNVs.

We have shown that variation in the cellular composition of intact human brain samples drives covariation of transcripts that are uniquely or predominantly expressed in specific kinds of cells^31–34^. We have also shown that the correlation between a gene’s expression pattern and the abundance of a cell type is a proxy for the extent to which the same gene is differentially expressed by that cell type^31^. These findings suggest that the core molecular properties of malignant clones can be identified in bulk tumor samples by correlating molecular feature levels with clonal abundance, which can be quantified through integrative analysis of variant allele frequencies (VAFs)^35,36^. In principle, such findings should be highly robust since they derive from millions or even billions of cells. Similar logic extends to nonmalignant cell types of the TME^31^.

Here we describe a novel strategy for deconstructing ITH through multiomic and multiscale analysis (MOMA) of serial tumor sections. By analyzing gene expression, whole exomes, deeply sequenced PCR amplicons spanning mutation sites, DNA methylation, and single-nucleus DNA and RNA, we exhaustively analyze ITH in IDH-mutant astrocytomas. Through integrative analysis of single-nucleotide variants (SNVs) and CNVs, we precisely define the evolutionary histories and spatial distributions of malignant clones. By comparing these distributions to gene expression data derived from the same tumor sections, we reveal clone-specific transcriptional profiles and validate them orthogonally through comparisons with normal human brain and single-nucleus analysis. Our findings suggest that a core set of genes is consistently expressed by the truncal clone of human astrocytomas, suggesting new therapeutic targets and a generalizable strategy for precisely deconstructing ITH and clarifying the core molecular properties of distinct cellular populations in solid tumors.

## Results

### Overview of MOMA

**Fig. 1a** depicts a heterogeneous human brain tumor specimen consisting of distinct malignant clones and nonmalignant cell types of the TME. By amplifying this specimen into a series of standardized biological replicates through serial sectioning, we introduce variation in cellular composition across sections (**Fig. 1b**), which are analyzed using multiscale (bulk and single-nucleus) and multiomic assays (**Fig. 1c**). Correlative analysis of cellular frequencies and molecular feature levels (e.g., gene expression levels) in bulk sections predicts optimal markers of distinct malignant clones and nonmalignant cell types (**Fig. 1d**), which are validated by single-nucleus analysis of interpolated sections and histology (**Fig. 1e**). MOMA therefore combines the power of bulk sampling with the precision of single-cell analysis to achieve the best of both worlds.

**Figure 1.**
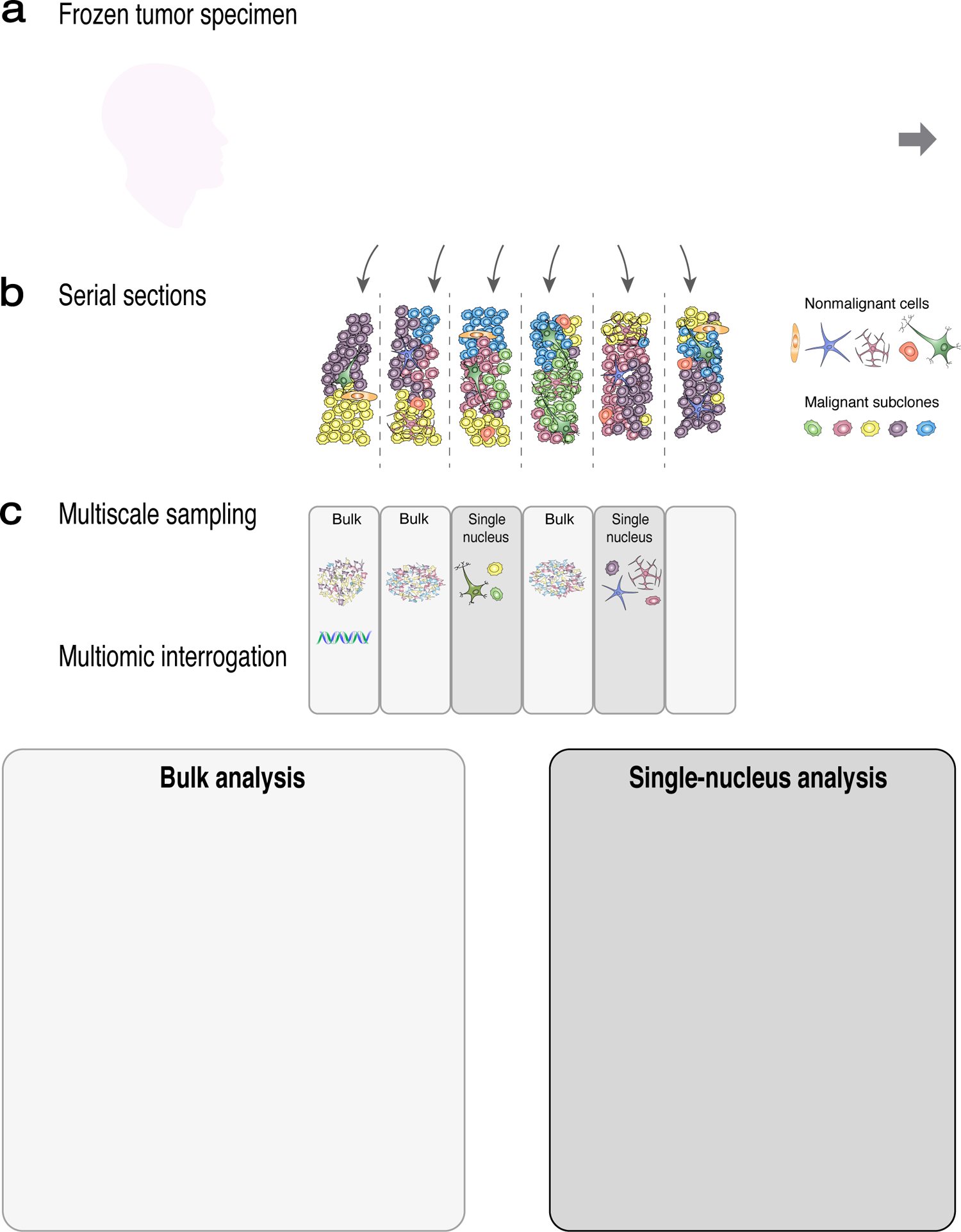
Overview of MOMA. **a)** Schematic of a heterogeneous human brain tumor. **b)** Serial sectioning introduces variation in cellular composition. **c)** Section usage can be flexibly tailored for diverse multiscale and multiomic assays. **d)** Correlative analysis of bulk cellular frequencies and molecular feature levels predicts optimal markers of malignant subclones and nonmalignant cell types of the tumor microenvironment. **e)** Predictions from bulk analysis are validated by single-nucleus analysis of interpolated sections and histology.

### Case 1: analysis of clonal composition

To put these ideas into practice, we obtained a resected specimen from a primary diffuse glioma that was removed from the left cerebral hemisphere of a 40 y.o. female who presented with language deficits (**Fig. 2a-c**). Molecular pathology revealed evidence for mutations in *IDH1* and *TP53* (**Fig. 2d-e**), no evidence for chromosome 1p/19q codeletion (data not shown), and KI67 labeling of 6% (data not shown), consistent with a CNS WHO grade 2 astrocytoma, IDH-mutant.

**Figure 2.**
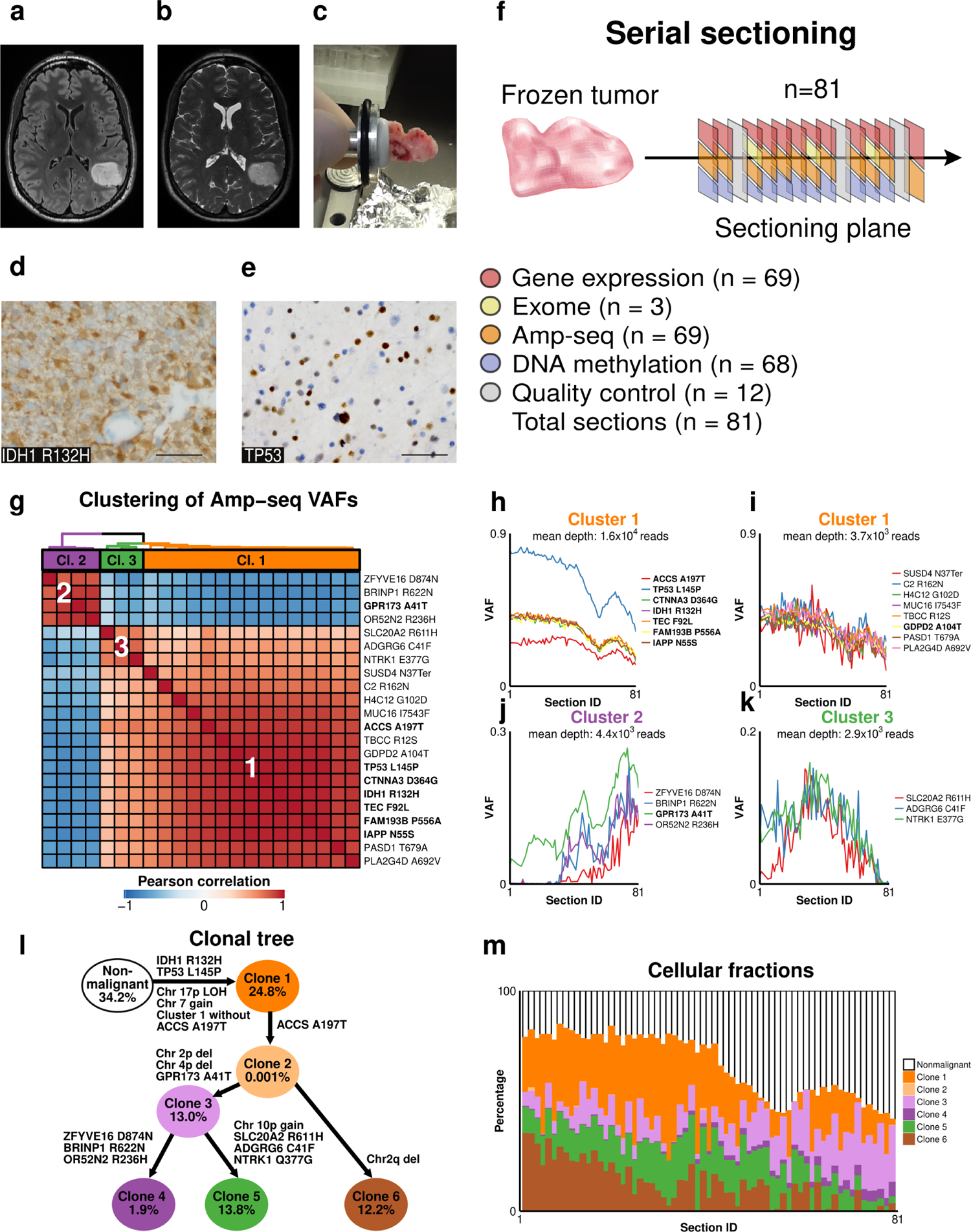
Multiomic analysis of serial tumor sections reveals the clonal composition of a primary grade 2 IDH-mutant astrocytoma (case 1). Axial T2 **(a)** and axial FLAIR **(b)** images demonstrate a round, well-defined T2 and FLAIR hyperintense intraaxial left temporoparietal mass that is non-enhancing and consistent with a low-grade glial neoplasm. **c)** Image of the frozen tumor sample prior to cryosectioning and nucleic acid isolation. **d-e)** Immunostaining for IDH1 R132H (**d**) and TP53 (**e**). Images: 400x. Scale bars: 50 μm. **f)** Schematic of serial sectioning strategy and section usage plan. Amp-seq = deep sequencing of PCR amplicons spanning mutations identified by exome sequencing. **g)** Hierarchical clustering of mutations, using 1 – Pearson correlation of amp-seq variant allele frequencies (VAFs) over all tumor sections (n = 69) as a distance measure, reveals three clusters. Amp-seq was performed in two sequencing runs (denoted by bold and regular fonts). **h-k)** VAF patterns comprising cluster 1 **(h,i)**, cluster 2 **(j)**, and cluster 3 **(k)**. Cluster 1 was split to illustrate the effects of high **(h)** and low **(i)** coverage. **l)** Clone phylogeny (with arbitrary branch lengths) derived from integrated analysis of SNVs (from amp-seq data) and CNVs (from DNA methylation data). Percentages represent the average abundance of each cellular fraction over all analyzed sections (n = 68). **m)** Estimated cellular fractions for all clones and nonmalignant cells over all sections (n = 68).

We cut 81 cryosections along the tumor specimen’s longest axis (**Fig. 2f**), followed by automated DNA/RNA extraction from each section (**Table S1**). To identify mutations and characterize the clonal landscape, we performed whole-exome sequencing (WES) on DNA isolated from sections 14, 39, 69, and the patient’s blood. Mutations detected in blood or in genes with very low tumor expression levels were excluded. Of the remaining 33 mutations (**Table S2**), including an in-frame deletion in *ATRX*, which is often mutated in IDH-mutant astrocytomas^37^, 18 were validated by Sanger sequencing and deep sequencing of PCR amplicons spanning each mutation (amp-seq; **Table S3**), five were validated only by amp-seq, and ten (mostly indels) could not be validated (**Table S2**). Among 22 validated coding mutations, 16 were detected by WES in all three tumor sections and six were detected in only one section, suggesting clonal heterogeneity among malignant cells (**Fig. S1a**).

To determine the relative abundance and spatial distributions of cells carrying mutations within the tumor specimen, we quantified VAFs for validated somatic mutations in all tumor sections using amp-seq (**Fig. S1b**). Amp-seq was performed in two sequencing runs: an initial run consisting of 25 amplicons (mean coverage: 3.0×10^3^ reads/mutation/section) and a second run consisting of nine amplicons (mean coverage: 1.7×10^4^ reads/mutation/section), with theoretical VAF detection sensitivity of <1%. To analyze the stability of amp-seq-derived VAFs, we downsampled reads spanning IDH1 R132H or TP53 L145P and calculated the root-mean-square-error (RMSE) and Pearson correlation between VAFs from full and downsampled read depths. This analysis revealed monotonic improvement in VAF estimates as a function of read depth (**Fig. S1c-d**). Importantly, VAFs derived from 100-200x coverage were far noisier than VAFs derived from full coverage, indicating that conventional WES data are inadequate for precisely estimating VAFs and malignant cell abundance.

We performed unsupervised hierarchical clustering of amp-seq data to identify mutations with similar VAF patterns within the tumor sample (**Fig. 2g** and **Table S4**). This analysis revealed three distinct clusters. Cluster 1 included 15 mutations with VAFs that decreased in the latter sections of the tumor sample, which were separated according to sequencing run to display the effects of read depth (**Fig. 2h,i**). Cluster 2 included four mutations with VAFs that increased in the latter sections of the tumor sample (**Fig. 2j**). Cluster 3 included three mutations with VAFs that peaked in the middle sections of the tumor sample (**Fig. 2k**).

Focusing on the sequencing run with higher coverage, we observed that five mutations in cluster 1 (including IDH1 R132H) had VAFs over all tumor sections that were statistically indistinguishable (**Fig. 2h**). Two other mutations (TP53 L145P and ACCS A197T) followed a similar pattern but at different scales. qPCR revealed approximately disomic copy numbers for both genes in all analyzed sections (**Fig. S1e**). These results indicate that VAFs for TP53 L145P reflect copy-neutral loss of heterozygosity for chromosome 17p (chr17p LOH) that occurred early in the tumor’s evolution (but after the L145P point mutation). Notably, the frequencies of chr17p LOH (derived from B-allele frequencies) were highly concordant between WES and amp-seq data (r=0.99, **Fig. S1f** [top]). In contrast, the lower VAFs for ACCS A197T suggest that this mutation appeared after the other mutations comprising cluster 1.

To determine the clonal composition and evolutionary history of the tumor specimen more precisely, we analyzed genome-wide CNVs and their relationships to SNVs quantified by amp-seq. CNVs were called from WES (n=3 sections) and DNA methylation (n=68 sections) data using FACETS^38^ and ChAMPS^39^, respectively, yielding highly concordant frequencies for copy number changes (r=0.92, **Fig. S1f** [bottom] and **Table S5**). Through combined analysis of SNV and CNV frequencies over all tumor sections, we produced an integrated model of tumor evolution. Specifically, we used PyClone^35^ to jointly analyze SNV and CNV frequencies, which identified seven distinct clusters and their overall prevalence. Subsequently, the evolutionary history of the tumor specimen was reconstructed using CITUP^36^, which produced the most likely phylogenetic tree (**Fig. 2l**) and frequencies of six malignant clones over all sections (**Fig. 2m** and **Table S6**). These analyses confirmed the truncal nature of mutations in *IDH1* and *TP53*^37^, while revealing wide variation in the purity of individual tumor sections (range: 38.3 - 84.8%; **Table S6**).

### Case 1: analysis of gene expression

We next explored relationships between clonal abundance and gene expression. We performed genome-wide gene coexpression analysis to identify groups of genes with similar expression patterns over all tumor sections. We identified 38 modules of coexpressed genes (arbitrarily labeled by colors), which were summarized by their eigengenes (i.e., first principal components) and hierarchically clustered (**Table S7, Fig. 3a-c**). As we have shown previously^31–34^, many modules were significantly enriched with markers of nonmalignant cell types (**Fig. S2a-d**). By comparing cumulative clonal abundance (**Fig. 2m**) to module eigengenes over all tumor sections, we identified five gene coexpression modules whose expression patterns closely tracked the abundance of clone 1 (turquoise: r=0.97, **Fig. 3d**), clone 3 (blue: r=0.84, **Fig. 3e**), clone 4 (black: r=0.83, **Fig. 3f**), clone 5 (midnightblue: r=0.71, **Fig. 3g**), and clone 6 (steelblue: r=0.69, data not shown). We did not identify a module that was significantly correlated with clone 2, which represented only 0.001% of cells (**Fig. 2l**).

**Figure 3.**
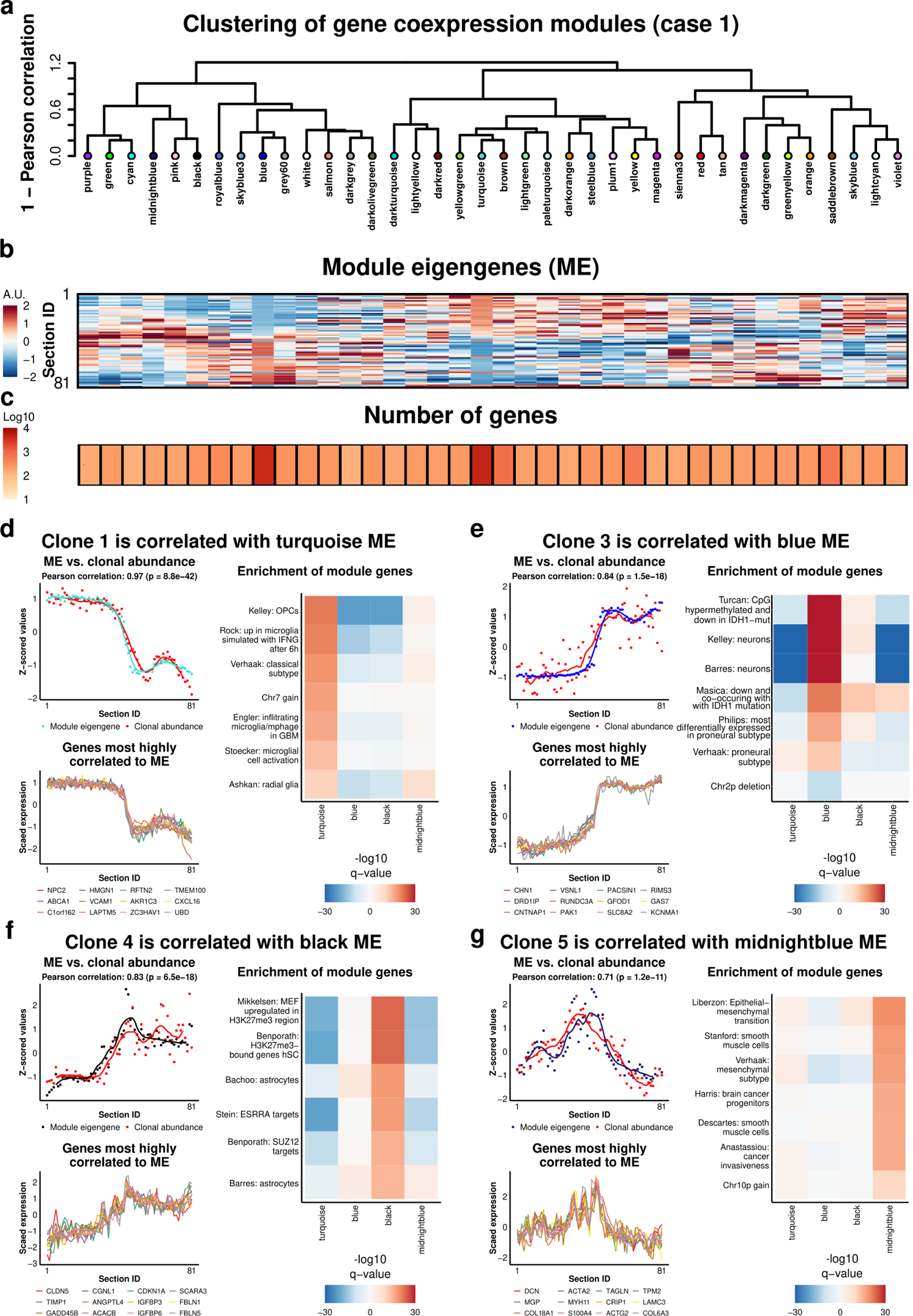
Gene coexpression modules are highly correlated with clonal abundance (case 1). **a)** Hierarchical clustering of gene coexpression modules over all tumor sections (n = 69). **b)** Module eigengenes (ME) illustrate the relative expression levels of genes in each module over all tumor sections. **c)** The number of genes used to form each ME. **d-g)** Top left: MEs with the strongest correlations to clonal abundance (defined cumulatively). Locally weighted smoothing (LOESS) lines are shown; correlation is based on data points. Bottom left: the 12 genes with the highest correlations to the ME (*k*_ME_). Right: enrichment analysis of gene coexpression modules using published gene sets. FDR-corrected p-values (q-values) from one-sided Fisher’s exact tests are shown. Positive values represent enrichments of genes that were significantly positively correlated to the ME, while negative values represent enrichments of genes that were significantly negatively correlated to the ME. Gene sets representing chromosomal gains or losses include all genes within affected regions (as described in Fig. 2l and **Table S5**). See **Table S9** for descriptions and sources of featured gene sets.

To characterize these modules, we performed enrichment analysis with biologically relevant gene sets (**Fig. 3d-g**). We first asked whether genes within clonal CNV boundaries (**Fig. 2l** and **Table S5**) were significantly enriched (for gains) or depleted (for deletions) in the bulk coexpression modules most strongly associated with each clone (**Table S8**). Notably, all such gene sets were significantly enriched in the appropriate module and expected direction (e.g., chr7 gain for clone 1 [**Fig. 3d**], chr2p deletion for clone 3 [**Fig. 3e**], and chr10p gain for clone 5 [**Fig. 3g**]). We next analyzed publicly available gene sets from diverse sources (**Table S9**). We found that the largest (turquoise) module, which closely tracked the abundance of clone 1 (i.e., tumor purity), was significantly enriched with markers of oligodendrocyte progenitor cells (OPCs) and radial glia, genes comprising the ‘classical’ subtype of glioblastoma proposed by Verhaak et al.^40^ and numerous gene sets related to microglial infiltration and activation. The second largest (blue) module, which tracked clone 3, was significantly enriched with neuronal gene sets as well as genes that are down-regulated pursuant to *IDH1* mutations. The black module, which tracked clone 4, was enriched with astrocyte markers as well as genes that are differentially regulated during development and glioma. The midnightblue module, which tracked clone 5, was enriched with markers of smooth muscle cells, genes comprising the ‘mesenchymal’ subtype of glioblastoma^40,41^, and gene sets related to epithelial-mesenchymal transition and invasiveness. The steelblue module, which tracked clone 6, was enriched with markers of non-resident immune cells (data not shown).

To further characterize the transcriptional signatures associated with each clone, we used multiple linear regression to model genome-wide expression levels as a function of clonal abundance. To account for collinearity and the dominant effect of clone 1, we used a group lasso model with bootstrapped clonal abundance vectors (real or permuted) as predictors (**Fig. S2e-i**). We restricted our focus to genes that were significantly and stably modeled by a single clone (in addition to clone 1, per the group lasso model, **Table S10**). Enrichment analysis of these genes largely recapitulated enrichment analysis of gene coexpression modules associated with each clone, including the associations of different clones with different cell types (**Table S11** and **Fig. S2i**).

The associations of different clones with different cell types suggest two non-mutually exclusive possibilities. First, different clones may preferentially express different cell-type-specific transcriptional programs. Second, different clones may preferentially associate with different nonmalignant cell types in the TME, leading to correlated gene expression patterns. Although such possibilities are ideally studied at the level of individual cells, all sections from this case were consumed during bulk data production. However, we reasoned that bona fide transcriptional signatures of malignant clones should be absent from non-neoplastic human brains. To test this hypothesis, we profiled gene expression in 361 cryosections from four neurotypical adult human brain samples (**Table S12**) and performed genome-wide differential coexpression analysis by subtracting normal correlations from tumor correlations, such that tumor-specific gene coexpression relations would be retained (**Fig. S3a, Table S13, Table S14**). This analysis revealed tumor-specific gene coexpression modules that tracked the abundance of distinct clones and largely recapitulated the transcriptional signatures described in **Fig. 3** and **Fig. S2**, including preserved enrichment of clone-specific CNV gene sets (**Fig. S3b-e**). However, enrichment results for nonmalignant cell-type-specific gene sets became less significant, with the exception of OPCs and radial glia for clone 1, which became more significant (**Fig. S3b-e**). These results suggest that derived clones may occupy distinct microenvironments, while the truncal clone retains signatures of progenitor cells that may reflect the cell of origin.

### Case 2: analysis of clonal composition

To test our strategy on a more complex case, we obtained a resected specimen from a recurrent diffuse glioma that was removed from the right cerebral hemisphere of a 58 y.o. male (**Fig. 4a-c**) approximately 28 years after the primary resection. Molecular pathology revealed evidence for mutations in *IDH1* and *TP53* (**Fig. 4d-e**), no evidence for chromosome 1p/19q codeletion (data not shown), and KI67 labeling of 4% (data not shown), consistent with a recurrent CNS WHO grade 2 astrocytoma, IDH-mutant. Building on our observations from case 1, we applied the same strategy to case 2, with five modifications. First, we increased power by analyzing more sections (**Table S15**). Second, we rotated the sample 90° halfway through sectioning to capture ITH in orthogonal planes (**Fig. 4f**). Third, we inferred CNVs from RNA-seq data instead of DNA methylation data. Fourth, we increased the average sequencing depth for amp-seq data. And fifth, we analyzed single nuclei from interpolated sections to validate predictions from bulk sections (**Fig. 4f**).

**Figure 4.**
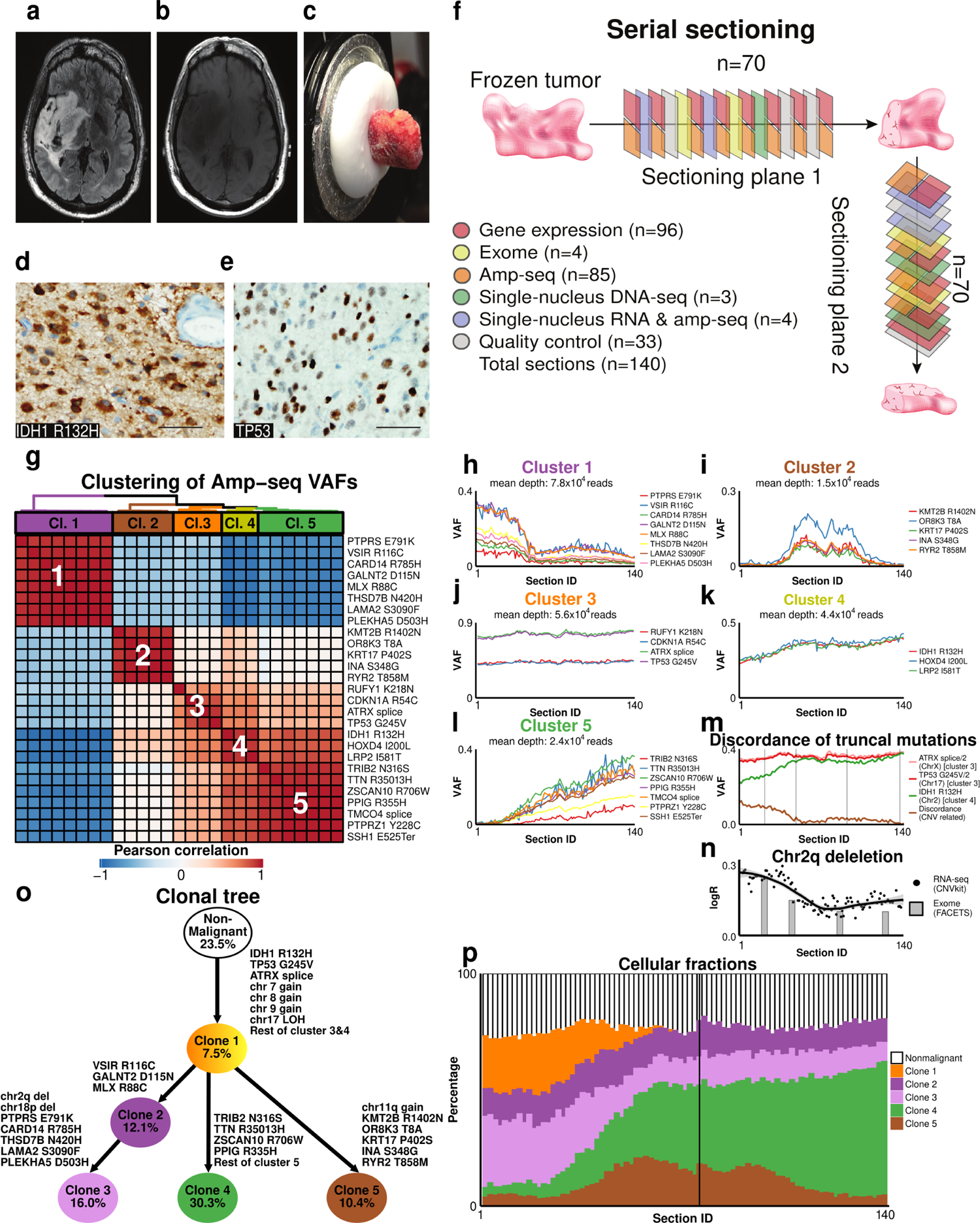
Multiomic analysis of serial tumor sections reveals the clonal composition of a recurrent grade 2 IDH-mutant astrocytoma (case 2). Axial T2 **(a)** and axial FLAIR **(b)** images demonstrate a non-enhancing, expansile, infiltrating glioma centered in the right insula and involving the basal ganglia, inferior frontal lobe, and temporal lobe. Cystic degeneration was present in the tumor. **c)** Image of the frozen tumor specimen prior to cryosectioning and nucleic acid isolation. **d)** The tumor was determined to harbor the IDH1 R132H mutation based on immunostaining with an antibody specific to the mutant protein. **e)** TP53 immunostaining demonstrated nuclear expression with an estimated staining index of 20%. All histological images were captured at 400x. Scale bars denote 50 μm. **f)** Schematic of serial sectioning strategy and section usage plan. **g)** Hierarchical clustering of mutations, using 1 – Pearson correlation of amp-seq VAFs over all tumor sections (n = 85) as a distance measure, reveals five clusters. **h-l)** VAF patterns comprising cluster 1 **(h)**, cluster 2 **(i)**, cluster 3 **(j)**, cluster 4 **(k)**, and cluster 5 **(l)**. **m)** Controlling for gene dosage reveals discordance of IDH1 R132H VAF with respect to truncal *ATRX* and *TP53* mutations, which is explained by a subclonal deletion of chromosome 2q (including *IDH1*) that occurred after the *IDH1* point mutation. **(n)** Heatmap of the chromosome 2q deletion event frequency (as determined by FACETS^38^), with LOESS fit line (black) and smoothed 95% confidence interval (gray envelope). **o)** Clone phylogeny (with arbitrary branch lengths) derived from integrated analysis of SNVs (from amp-seq data) and CNVs (from RNA-seq data). Percentages represent the average abundance of each cellular fraction over all analyzed sections (n = 85). **p)** Estimated cellular fractions for all clones and nonmalignant cells over all sections. Black vertical line denotes orthogonal sample rotation.

To identify somatic mutations, we performed WES on DNA from two sections in each plane (22, 46, 85, 123; **Table S16**) and the patient’s blood. 227 mutations were identified and 74 were selected for amp-seq by clustering WES VAFs to reveal candidate mutations most likely to mark distinct clones (**Table S17**). Of these, 58 mutations were verified by amp-seq (**Table S18**).

As with case 1, downsampling reads spanning IDH1 R132H or TP53 G245V revealed monotonic improvements in VAF estimates as a function of read depth (**Fig. S4a-b**). We therefore restricted further analysis of amp-seq data to 27 mutations with high coverage over all tumor sections or strong VAF correlations to other mutations (**Fig. S4c**). Hierarchical clustering of these amp-seq data (**Table S18**) revealed five clusters of mutations with similar VAF patterns within the tumor sample (**Fig. 4g-l**), suggesting multiple malignant clones.

Because mutations in *IDH1*, *TP53*, and *ATRX* are considered diagnostic for astrocytoma^37^, we expected these to be truncal and were therefore surprised that IDH1 R132H fell in a separate cluster from mutations in *TP53* and *ATRX* (**Fig. 4j-k**). To explore this discrepancy, we analyzed VAFs for all three mutations after controlling for gene dosage. This analysis revealed greater discordance between VAFs for *IDH1* and *TP53* / *ATRX* mutations in sectioning plane 1 vs. sectioning plane 2 (**Fig. 4m**). We also observed that all genes in mutation cluster 4 (including *IDH1*) are located on chr2q. These observations suggested that the discrepancy between *IDH1* and *TP53* / *ATRX* mutation VAFs might be explained by a subclonal deletion in chr2q pursuant to the IDH1 R132H mutation, as has been previously reported^42–44^. To test this hypothesis, we quantified CNVs from WES (n=4 sections) and RNA-seq (n=90 sections) data using FACETS^38^ and CNVkit^45^, respectively, which yielded highly concordant frequencies for copy number changes (r=0.97, **Fig. S4d** and **Table S19**), including a chr2q deletion event. As expected, frequencies of the chr2q deletion event were substantially higher in sectioning plane 1 vs. sectioning plane 2 (**Fig. 4n**) and almost perfectly correlated with the observed discordance between *IDH1* and *TP53* / *ATRX* mutation VAFs (r=0.98, **Fig. S4e**).

Through combined analysis of SNV and CNV frequencies over all tumor sections, we generated an integrated model of tumor evolution using the same approach described for case 1, including the most likely phylogenetic tree (**Fig. 4o**) and frequencies of five malignant clones over all sections (**Fig. 4p** and **Table S20**). Compared to case 1, there was substantially less variation in the purity of individual tumor sections (range: 71.4 - 81.6%; **Table S20**). We confirmed the truncal nature of mutations in *IDH1*, *TP53*, and *ATRX*, along with gains of chr7, chr8, and chr9. To more closely examine the sequence of early mutational events, we performed single-nucleus DNA sequencing using MissionBio’s Tapestri microfluidics platform^46^. We took advantage of an existing panel of cancer genes, which included primers flanking one *IDH1* and two *TP53* loci. We were also able to infer chr17 and chr2q copy-number changes using mutations that fell within the targeting panel. We analyzed 4,433 nuclei from plane 1 (section 29) and 3,736 nuclei from plane 2 (sections 113 and 115). Clustering nuclei from each plane revealed clonal frequencies that broadly matched those obtained by bulk analysis (**Fig. S4f, Table S21**). Interestingly, we observed a subpopulation of clone 1 (clone 1a: 4.1 - 6.6%) with IDH1 R132H -/+ and TP53 G245V -/+/+ genotypes (**Fig. S4f**). These genotypes suggest that *TP53* LOH occurred mechanistically in this case through duplication of the mutant allele prior to loss of the wild-type allele, and may also explain the slightly lower VAFs for TP53 G245V compared to the mutation in *ATRX* (**Fig. 4j**).

### Case 2: analysis of gene expression

We explored relationships between clonal abundance and gene expression using the same strategies described for case one. Genome-wide gene coexpression analysis identified 68 modules of coexpressed genes, which were summarized by their eigengenes and hierarchically clustered (**Fig. 5a-c**). As expected^31,32^, many modules were significantly enriched with markers of nonmalignant cell types (**Fig. S5a-d**). By comparing clonal abundance (**Fig. 4p, Table S20**) to module eigengenes over all tumor sections, we identified five gene coexpression modules whose expression patterns closely tracked the abundance of clone 1 (red: r=0.65, **Fig. 5d**), clone 2 (violet: r=0.82, **Fig. 5e**), clone 3 (black: r=0.8, **Fig. 5f**), clone 4 (ivory: r=0.86, **Fig. 5g**), and clone 5 (lightcyan: r=0.82, data not shown).

**Figure 5.**
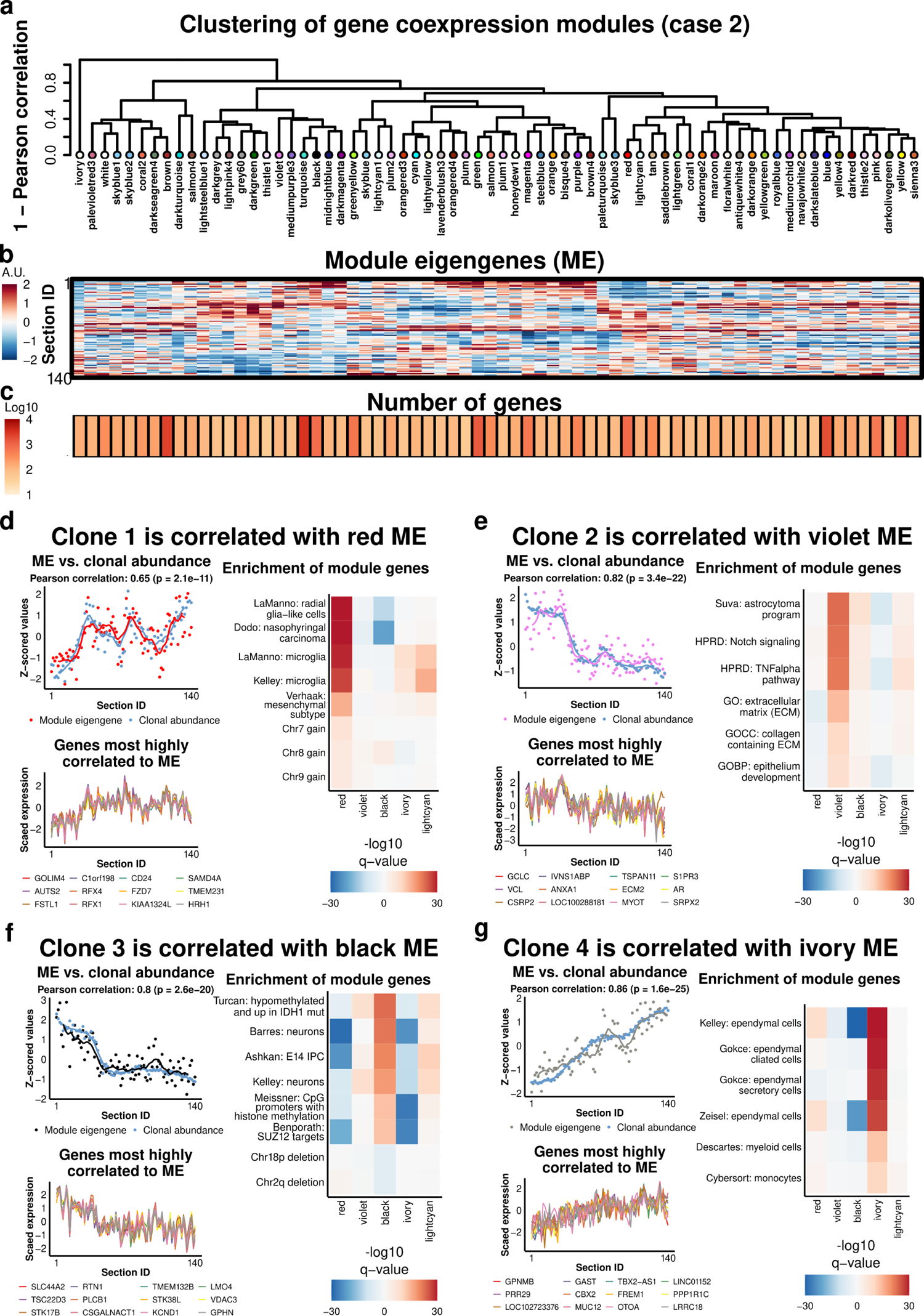
Gene coexpression modules are highly correlated with clonal abundance (case 2). **a)** Hierarchical clustering of gene coexpression modules over all tumor sections (n = 90). **b)** Module eigengenes (ME) illustrate the relative expression levels of genes in each module over all tumor sections. **c)** The number of genes that formed each ME. **d-g)** Top left: MEs with the strongest correlations to clonal abundance (defined cumulatively). Locally weighted smoothing (LOESS) lines are shown; correlation is based on data points. Bottom left: the 12 genes with the highest correlations to the ME (*k*_ME_). Right: enrichment analysis of gene coexpression modules using published gene sets. FDR-corrected p-values (q-values) from one-sided Fisher’s exact tests are shown. Positive values represent enrichments of genes that were significantly positively correlated to the ME, while negative values represent enrichments of genes that were significantly negatively correlated to the ME. Gene sets representing chromosomal gains or losses include all genes within affected regions (as described in Fig. 4o and **Table S19**). See **Table S9** for descriptions and sources of featured gene sets.

Enrichment analysis using gene sets defined by clonal CNV boundaries (**Fig. 4o** and **Table S19**) confirmed expected over-representation (for gains) or under-representation (for deletions) in the bulk coexpression modules most strongly associated with each clone (**Fig. 5d-g, Table S22, Table S23)**. Further analysis using publicly available gene sets from diverse sources (**Table S9**) revealed that the red module, which tracked the abundance of clone 1 (i.e., tumor purity), was significantly enriched with markers of radial glia and microglia, as well as genes comprising the mesenchymal subtype of glioblastoma. The violet module, which closely tracked the abundance of clone 2, was significantly enriched with genes from reported astrocytoma expression programs, as well as TNFalpha signaling and extracellular matrix components. The black module, which closely tracked the abundance of clone 3, was significantly enriched with markers of neurons and genes involved in chromatin remodeling. The ivory module, which closely tracked the abundance of clone 4, was enriched with markers of ependymal cells and myeloid cells. The lightcyan module, which closely tracked the abundance of clone 5, was significantly enriched with genes involved in EGFR and NF-kB signaling, as well as genes comprising the proneural subtype of glioblastoma (data not shown).

To further characterize the transcriptional signatures associated with each clone, we used multiple linear regression to model genome-wide expression levels as a function of clonal abundance. To account for collinearity, we used a regular lasso model with bootstrapped clonal abundance vectors (real or permuted) as predictors (**Fig. S5e-i**). We restricted our focus to genes that were significantly and stably modeled by a single clone (**Table S24)**. Enrichment analysis of these genes largely recapitulated enrichment analysis of gene coexpression modules associated with each clone, including CNVs and the associations of different clones with different cell types (**Fig. S5i, Table S25**).

To validate gene expression signatures of malignant clones and nonmalignant cell types identified from bulk tumor sections, we performed single-nucleus RNA-seq (snRNA-seq) on tumor sections 17, 53, 93, and 117 (**Fig. 4f, Table S26**). Using a protocol adapted from TARGET-Seq^47,48^, we profiled gene expression in 288 flow-sorted nuclei per section. Following data preprocessing and quality control, 809 nuclei (70.2%) with an average of >200K unique reads/nucleus were retained for further analysis. Uniform manifold approximation and projection (UMAP) analysis revealed that nuclei did not segregate by section ID (**Fig. S6a, Table S27**).

To determine whether nuclei segregated by cancerous state, we analyzed the malignancy of each nucleus. Unlike some tumors, astrocytomas are not defined by truncal CNVs, which can drive gene expression changes that are used to infer malignancy in snRNA-seq data^25,37,49,50^. We therefore genotyped all nuclei through single-nucleus amplicon sequencing (snAmp-seq) of cDNA spanning mutations in the truncal clone (**Fig. 4o**). This analysis provided sufficient information to call malignancy for 75% of nuclei. Projecting malignancy status onto the UMAP plot revealed clear segregation of malignant and nonmalignant nuclei (**Fig. S6b**).

To further classify nuclei as specific malignant clones or nonmalignant cell types, we took a two-step approach. First, we hierarchically clustered all nuclei using a Bayesian distance metric calculated by Sanity^27^ that downweights genes with large error bars, revealing 12 clusters. Second, we asked whether genes in the bulk coexpression modules most strongly associated with each malignant clone or nonmalignant cell type were upregulated in distinct snRNA-seq clusters compared to all other genes (**Fig. S7a-j**). This analysis revealed specific and significant upregulation of genes from the red (**Fig. 5d**), violet (**Fig. 5e**), black (**Fig. 5f**), and lightcyan (data not shown) modules in snRNA-seq clusters 2, 1, 7, and 10 (**Fig. 6a**), suggesting that these clusters correspond to malignant clones 1, 2, 3, and 5, respectively. Genes in the ivory module (**Fig. 5g**) were significantly upregulated in snRNA-seq clusters 3 and 5, suggesting that both of these clusters represent clone 4 (**Fig. 6a**). Similarly, we observed specific and significant upregulation of genes from the purple (**Fig. S5a**), yellow (**Fig. S5c**), green (**Fig. S5d**), and orange (data not shown) modules in snRNA-seq clusters 9, 4, 12, and 6 (**Fig. 6a**), suggesting that these clusters correspond to nonmalignant astrocytes, microglia, neurons, and endothelial cells, respectively. Genes in the tan module (**Fig. S5b**) were significantly upregulated in snRNA-seq clusters 8 and 11, suggesting that both of these clusters represent nonmalignant oligodendrocytes (**Fig. 6a**).

**Figure 6.**
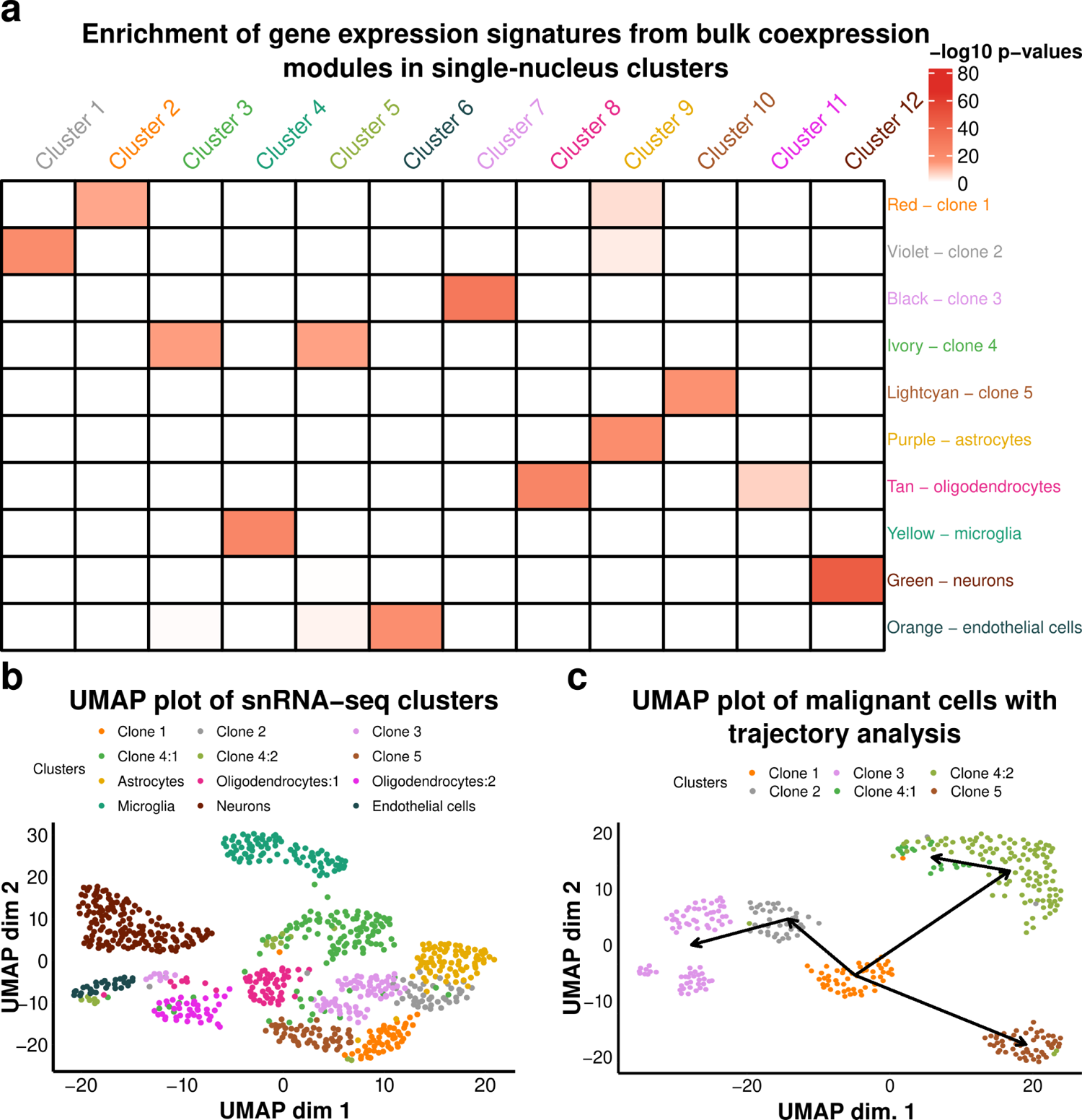
Single-nucleus RNA-seq analysis validates inferences from bulk data. **a)** Heatmap of P-values (one-sided Wilcoxon rank-sum test) comparing differential expression t-values for genes comprising each bulk coexpression module (colors, x-axis) to all other genes in each SN cluster versus all other clusters. **b)** UMAP plot of all nuclei (n = 809) with characterizations of clusters from (**a**) superimposed. **c)** UMAP plot of malignant nuclei (n = 360), with results of Slingshot trajectory analysis^51^ superimposed. Malignancy was determined by genotyping all nuclei via single-nucleus amplicon sequencing (snAmp-seq) of cDNA spanning mutations in the truncal clone.

We performed several additional analyses to verify these findings. First, we projected snRNA-seq cluster assignments onto the UMAP plot (**Fig. 6b**) and observed that cluster assignments were consistent with the malignancy map produced by genotyping nuclei via snAmp-seq (**Fig. S6b**). Second, we performed UMAP analysis for malignant cells only, followed by trajectory analysis with Slingshot^51^ (**Fig. 6c**). This analysis revealed patterns of clonal evolution that recapitulated the phylogenetic tree inferred from integrative analysis of bulk tumor sections (**Fig. 4o**). Third, we compared estimates of cellular abundance obtained from bulk and single-nucleus data for adjacent tissue sections. This analysis revealed highly consistent estimates for the relative abundance of malignant clones (r ≥ 0.94; **Fig. S6c**) and nonmalignant cell types (r ≥ 0.90; **Fig. S6d**).

Supervised clustering with differentially expressed genes revealed clear separation of snRNA-seq clusters (**Fig. 7**). Overall, malignant clones were more transcriptionally active than nonmalignant cell types, with the exceptions of clone 4:1 and endothelial cells (**Fig. 7**, right). Enrichment analysis of genes that were significantly up-regulated in snRNA-seq clusters confirmed the identities of nonmalignant cell types (**Fig. S8, Table S28**). For malignant clones, enrichment analysis of snRNA-seq clusters supported and refined inferences from bulk data (**Fig. 5d-g**, **Fig. S5i**, **Fig. S6**, **Table S9**). For clone 1, consistent enrichments for markers of radial glia and genes comprising the mesenchymal subtype of glioblastoma were observed in bulk and snRNA-seq data. In contrast, markers of microglia were less significantly enriched in clone 1 nuclei from snRNA-seq data versus bulk data, and markers of oligodendrocyte progenitor cells (OPCs) were more significantly enriched. For clone 2, markers of astrocytes were more significantly enriched in snRNA-seq data versus bulk data. Clone 3 was consistently enriched with genes involved in chromatin remodeling, but neuronal markers were less significantly enriched in snRNA-seq data. Clone 4 showed strong enrichment for markers of ependymal cells in all analyses, while clone 5 was significantly enriched with genes comprising the proneural subtype of glioblastoma in all analyses. Interestingly, genes involved in mitosis were most highly expressed by clone 1, clone 4:2, and endothelial cells (**Fig. 7**, right).

**Figure 7.**
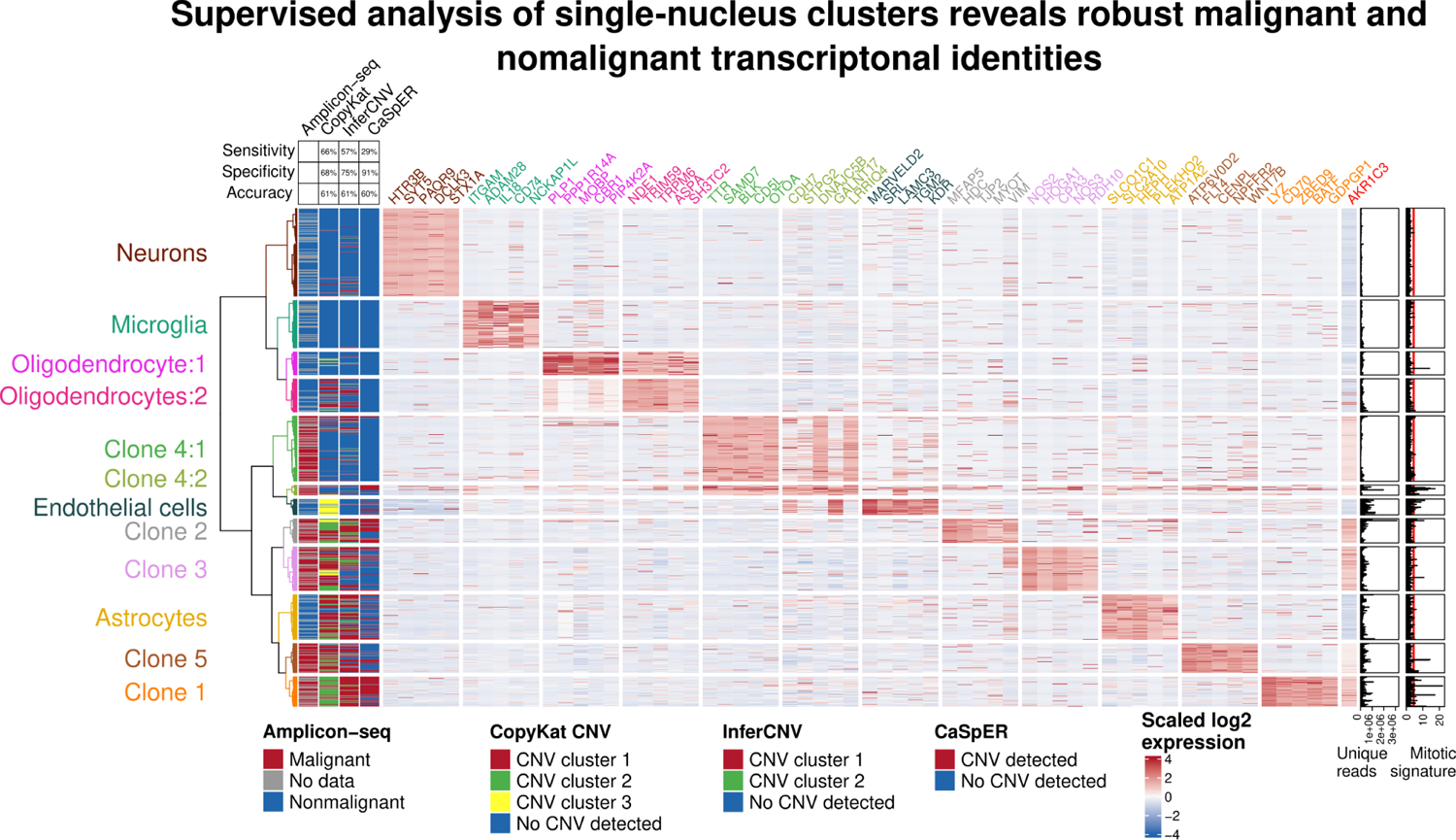
Genotyping nuclei profiled by snRNA-seq reveals the limitations of single-cell CNV-calling algorithms. Heatmap of scaled log_2_ expression vectors for the five most upregulated genes in each snRNA-seq cluster vs. all other clusters (one-sided Wilcoxon rank-sum test). Far left: malignancy vector determined by snAmp-seq of cDNA spanning mutations in the truncal clone. Left: malignancy vectors inferred from CNV analysis of snRNA-seq data using the CopyKat^49^, InferCNV^25^, or CaSpER^50^ algorithms (blue = nonmalignant; all other colors = malignant). Right: bar plots depict the total number of unique reads (UMIs) for each nucleus and the average number of UMIs for genes comprising the Gene Ontology category ‘mitotic chromosome condensation’ (GO: 0030261). Red vertical line: max expression of mitotic genes in neurons, which presumably represents background noise.

Because clones in this case were characterized by disparate CNVs (**Fig. 4o**), we asked how malignancy calls compared between algorithms that infer CNVs from snRNA-seq data and malignant genotypes derived from snAmp-seq data. We used CopyKat^49^, InferCNV^25^, and CaSpER^50^ to call CNVs from snRNA-seq data. These analyses revealed substantial variation in malignancy calls for different algorithms (**Fig. 7**) as well as differences from bulk CNV calls (e.g., no gains in chr7p, chr8p, and chr9q; **Fig. S6e**). Taking the snAmp-seq genotyping as ground truth, CopyKat and InferCNV were more sensitive but less specific than CaSpER, leading to discrepant calls. For example, nonmalignant astrocytes and oligodendrocytes:2 were mostly called malignant by CopyKat and InferCNV, while clone 4:2 was mostly called nonmalignant by these two algorithms. CaSpER’s classification of nuclei from these populations was mostly correct, but it failed to recognize most malignant nuclei for clones 3 and 5. In addition, clone 4:1 was mostly classified as nonmalignant by all three algorithms. Overall, none of the algorithms for inferring malignancy from CNVs achieved accuracy > 61% (**Fig. 7**).

### Integrative analysis of gene expression in malignant cells

Intuitively, genes whose expression patterns correlate most strongly with the abundance of malignant cells should include optimal biomarkers. This intuition can also be proven mathematically and empirically. **Fig. 8a-c** illustrates a hypothetical example in which the goal is to identify optimal transcriptional markers of malignant cells in a human brain tumor. A conventional strategy would involve physically isolating individual cells, transcriptionally profiling them by single-cell RNA-seq (scRNA-seq), inferring the malignancy of individual cells from the scRNA-seq data based on the presence of driver mutations (CNVs and/or SNVs), and performing differential expression analysis for each gene between all malignant and nonmalignant cells (for example, using a t-test; **Fig. 8b**). **Fig. 8c** shows an alternative analytical path that leads to the same place: by correlating expression levels of the same hypothetical gene from **Fig. 8b** with a dichotomous variable denoting malignant cell abundance (1=malignant cells, 0=nonmalignant cells), the resulting statistical significance is identical to that obtained by differential expression analysis.

**Figure 8.**
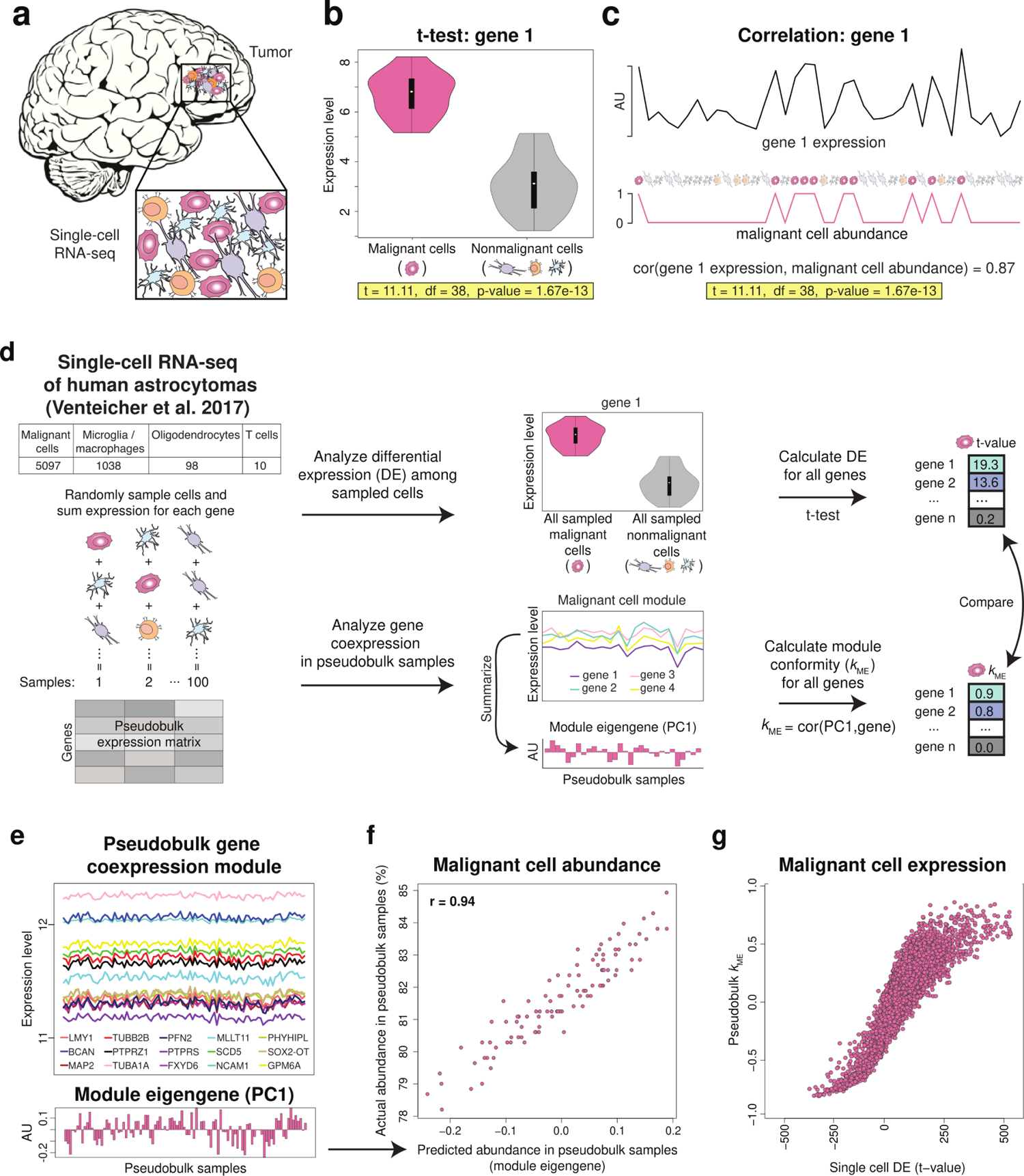
Correlation to malignant cell abundance predicts single-cell differential expression analysis of malignant vs. nonmalignant cells. **a-d)** Analysis schematic. An adult malignant glioma consisting of malignant cells (pink) interspersed with nonmalignant cells **(a)**. **b)** Single-cell RNA-seq (scRNA-seq) reveals a hypothetical gene (gene *X*) that is significantly up-regulated in malignant vs. nonmalignant cells. **c)** Correlating the same gene’s expression pattern with a binary vector encoding malignant cell abundance (1 = malignant, 0 = nonmalignant) produces identical results. **d)** Left: scRNA-seq data from 10 adult human IDH-mutant astrocytomas^24^ were randomly sampled and aggregated to create 100 pseudobulk samples. Right (top): Genome-wide differential expression (DE) was analyzed for all sampled cells. Right (bottom): Genome-wide gene coexpression was analyzed for all pseudobulk samples. Each pseudobulk module was summarized by its module eigengene (PC1), which was compared to malignant cell abundance, and the correlation between each gene and each module eigengene (module conformity, or *k*_ME_) was calculated. **e)** A pseudobulk malignant cell module featuring the top 15 genes ranked by *k*_ME_. By correlating the module eigengene to pseudobulk tumor purity **(f)**, we see that this module is driven by variation in malignant cell abundance among pseudobulk samples. **g)** The extent of DE (t-value) identified by scRNA-seq of malignant vs. nonmalignant cells predicts the correlation between gene expression and malignant cell abundance (pseudobulk *k*_ME_).

Although the t-test and correlation produce identical results when the independent variable is dichotomous, this is not the case when the independent variable is continuous. However, we have shown via pseudobulk analysis of scRNA-seq data from normal adult human brain that: i) the correlation between the expression pattern of a gene and the [continuous] abundance of a cell type accurately predicts differential expression of that gene in that cell type, and ii) cell-type-specific gene coexpression relationships accurately predict cellular abundance in pseudobulk samples^31^. To determine whether these findings extend to malignant cells, we repeated this analysis using scRNA-seq data from 10 adult human astrocytomas^24^ (**Fig. 8d**). Genome-wide gene coexpression analysis of pseudobulk samples obtained by randomly aggregating scRNA-seq data revealed a malignant cell coexpression module whose first principal component (or eigengene^52^) closely tracked the actual abundance of sampled malignant cells (**Fig. 8e-f**). Furthermore, the genes that were most significantly up-regulated in malignant cells per differential expression analysis of the underlying scRNA-seq data (**Fig. 8d**) also had the highest correlations to malignant cell abundance (*k*_ME_) in pseudobulk data (**Fig. 8g**). These results confirm that optimal markers of malignant cells can be revealed by correlating genome-wide expression patterns with malignant cell abundance in heterogeneous tumor samples. This strategy also applies to individual malignant clones (as well as nonmalignant cell types of the TME), as shown for case 2 in **Fig. S9**.

We next sought to compare transcriptional profiles of malignant cells between case one and case two through integrative analysis. However, despite the fact that both tumors were diagnosed as grade 2 IDH-mutant astrocytomas, only one SNV was shared between the cases. Furthermore, the shared SNV (IDH1 R132H) was absent in ∼21% of malignant cells in case 2 following loss of chr2q (**Fig. 4o**). We therefore asked whether the truncal clones (i.e., clone 1), which presumably included all of the mutations required to initiate these tumors along with passenger mutations, had consistent transcriptional profiles in case 1 and case 2. For each case, we analyzed genome-wide correlations to the cumulative abundance of clone 1 (equivalent to tumor purity). Comparing these results between cases, we observed a highly significant relationship (**Fig. 9a, Table S29**). Enrichment analysis of genes whose expression patterns were most positively correlated with clone 1 in both cases implicated gene sets comprising the ‘classical’ subtype of glioblastoma proposed by Verhaak et al.^40^, markers of radial glia, infiltrating monocytes, and extracellular matrix components (**Fig. 9a-b**; red). In contrast, genes whose expression patterns were most negatively correlated with clone 1 in both cases largely implicated gene sets related to neurons and neuronal function (**Fig. 9a-b**; blue).

**Figure 9.**
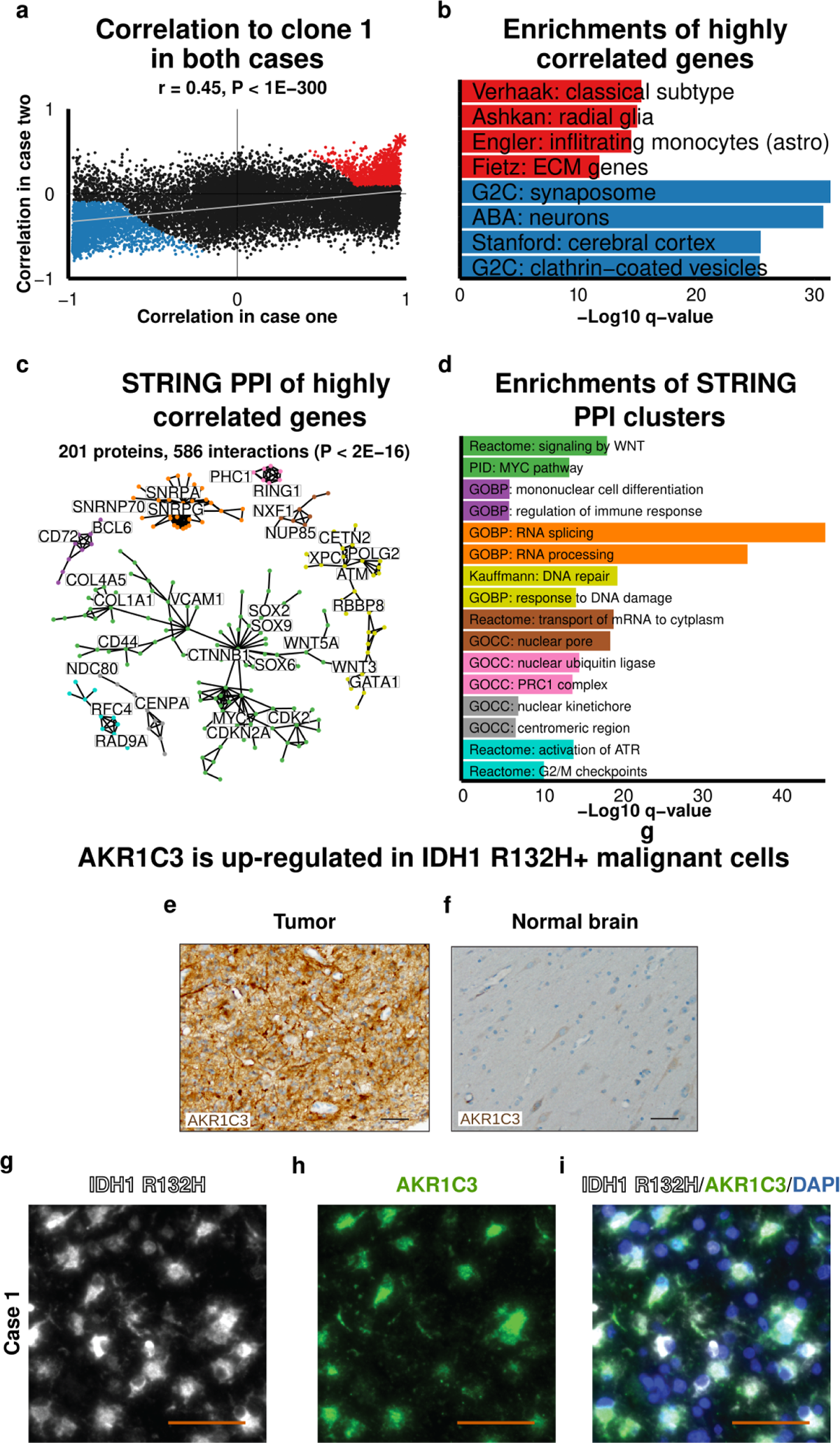
Aggregating correlations to tumor purity reveals core transcriptional features of astrocytomas. **a)** Gene expression correlations (n = 15,288 genes) to malignant cell abundance in case 1 and case 2. Red and blue denote significantly correlated genes that were used for enrichment analysis **(b)**, and the star denotes *AKR1C3*. **b)** -Log_10_ FDR-corrected p-values (q-values) from one-sided Fisher’s exact tests analyzing gene set enrichment in red and blue genes from **(a)**. **c)** Validated protein-protein interactions (PPI) from STRINGdb^54^ for red genes from **(a)**. The 201 proteins shown formed networks of five or more proteins, with the number of interactions equal to the number of edges. **d)** -Log_10_ FDR-corrected p-values (q-values) from one-sided Fisher’s exact tests analyzing gene set enrichment for each STRINGdb interaction cluster in **(c)**. **e-f)** AKR1C3 immunostaining in FFPE tissue adjacent to the sectioned region of case 1 **(e)** and non-neoplastic human brain **(f)**. Image: 200x; scale bar: 50 μm. **g-i)** Immunofluorescent co-staining of IDH1 R132H (white), AKR1C3 (green), and nuclei (blue [DAPI]) in case one demonstrating expression of AKR1C3 in malignant cells carrying the truncal IDH1 R132H mutation. Scale bar denotes 50µm.

We further characterized genes whose expression patterns were most positively correlated with the truncal clone in both cases (**Fig. 9a**; red) by cross-referencing them with human protein-protein interaction (PPI) data from the STRING database^53,54^. This analysis revealed eight distinct clusters of interacting proteins (**Fig. 9c**). The largest of these (green) included several SOX transcription factors and was significantly enriched with genes involved in WNT and MYC signaling (**Fig. 9d**). The second largest cluster (yellow) was significantly enriched with genes involved in DNA repair, and the third largest cluster (orange) was significantly enriched with genes involved in RNA splicing (**Fig. 9d**). The remaining clusters were significantly enriched with genes involved in mRNA transport (brown), DNA replication (turquoise), specific cellular compartments and protein complexes (pink, gray), and immune response (purple) (**Fig. 9d**).

To provide further validation for these findings, we performed immunostaining for AKR1C3. Out of 15,288 genes, *AKR1C3* bulk expression correlations to tumor purity ranked fifth in case one and first in case two (**Fig. 9a** [asterisk], **Table S29**). *AKR1C3* was also significantly upregulated in malignant vs. nonmalignant nuclei per snRNA-seq (**Fig. 7**, right). Immunostaining confirmed substantial upregulation of AKR1C3 in tumor vs. normal human brain at the protein level (**Fig. 9e**, **f**). To provide cellular resolution, we co-stained for AKR1C3 and IDH1 R132H using an antibody that recognizes the mutated IDH1 protein. As expected, this analysis revealed broad overlap between cells expressing AKR1C3 and cells expressing IDH1 R132H (**Fig. 9g-i**).

## Discussion

We have described a novel strategy called MOMA for deconstructing ITH through multiomic and multiscale analysis of serial tumor sections. By amplifying each tumor specimen into standardized biological replicates through serial sectioning, we obtained a large number of representative subsamples of each tumor with variable cellular composition. Because section size and number can be tailored to experimental needs, MOMA provides flexibility for a variety of concurrent assays while preserving spatial information. We performed WES to identify mutations in a small number of distant sections, followed by deep sequencing of PCR amplicons spanning mutation sites to quantify SNV frequencies with high confidence in a large number of sections. Although clusters of SNVs with highly correlated VAFs suggested distinct clones, integrative analysis of SNV and CNV frequencies (inferred from bulk DNA methylation data [case 1] or bulk RNA-seq data [case 2]) was required to accurately reconstruct clonal phylogenies. Using this approach, we identified the six most prevalent clonal populations of malignant cells in case 1 and five in case 2 and quantified their abundance in all tumor sections. By comparing clonal abundance to genome-wide expression patterns over all tumor sections, we identified transcriptional profiles of distinct malignant clones in each case. Clone expression profiles were orthogonally validated through comparisons with normal human brain (case 1) and snRNA-seq using nuclei isolated from interpolated tumor sections (case 2). Analysis of these profiles revealed several interesting findings. First, markers of neural stem cells (radial glia) were most significantly enriched in the truncal clone. Second, gene sets representing transcriptional subtypes of glioblastoma^40^ were significantly associated with distinct clones, suggesting stereotyped and potentially reversible malignant cell states that may reflect different microenvironments^55–57^. Third, single-nucleus expression signatures of mitosis were observed in most clones, suggesting they were not terminally differentiated. And fourth, genotyping nuclei analyzed by snRNA-seq revealed ependymal-like cells with oncogenic mutations, but no neuron-like cells with oncogenic mutations. To our knowledge, malignant ependymal cells have not previously been described in human astrocytomas. Because ependymal cells normally derive from radial glia during early brain development^58,59^, the presence of malignant ependymal cells (and the absence of malignant neurons) may point to a specific type of progenitor as the cell of origin for case two. Alternatively, oncogenic mutations and / or resulting epigenetic changes may alter the differentiation potential of the cell of origin from its normal state. In either case, replication of our strategy on malignant glioma subtypes will clarify whether brain tumors with specific oncogenic mutations consistently produce malignant progeny with transcriptional profiles that resemble the same neurobiological cell types.

Although both cases were diagnosed as IDH-mutant grade 2 astrocytomas, they shared only one SNV (IDH1 R132H), which was truncal in both cases but lost from 21% of malignant cells (clone 3) in case 2 due to chr2q deletion. The extent of clonal heterogeneity, even for the same type of tumor, begs the question of how gene expression correlations to clonal abundance should be compared and integrated across cases. We reasoned that aggregating gene expression correlations to the truncal clone (equivalent to tumor purity) would identify the most specific and consistent transcriptional features of all malignant cells in both astrocytomas (as illustrated in **Fig. 8**). This deceptively simple strategy has important implications for target discovery in cancer biology, because correlations between molecular abundance and tumor purity can be aggregated from huge numbers of bulk samples from similar cases that collectively represent many billions of cells. In statistical and economic terms, this strategy represents an efficient path for identifying robust molecular markers of malignant cells, including the non-oncogene dependencies that may vastly outnumber recurrently mutated genes^60^.

We observed a highly significant genome-wide correlation between gene expression profiles of the truncal clone, suggesting that a core set of genes is consistently expressed by the founding population of malignant cells in IDH-mutant astrocytomas. This result is particularly striking given the biological and technical differences between case 1 (primary astrocytoma, microarray gene expression data) and case 2 (recurrent astrocytoma, RNA-seq gene expression data). Cross-referencing these genes with human PPI data^54^ revealed distinct groups of interacting proteins that were significantly enriched with cancer-related pathways and processes, including WNT and MYC signaling, RNA splicing, and DNA repair. Furthermore, many of the genes whose expression patterns correlated most strongly with malignant cell abundance in both cases (**Table S29**) have been implicated in other types of cancer. For example, *AKR1C3*, which encodes a prostaglandin synthase involved in androgen production^61^, is significantly upregulated and associated with poor outcomes in hepatocellular carcinoma^62^, prostate cancer^63^, and pediatric T-cell acute lymphoblastic leukemia^64^. These findings underscore the possibility that malignant cells from diverse cancers caused by distinct mutations may nevertheless share transcriptional dependencies that can be exploited therapeutically.

It is also important to note that transcriptional phenotypes of malignancy, including upregulation of *AKR1C3*, persisted in clone 3 from case 2 despite loss of the driver mutation IDH1 R132H following chr2q deletion. IDH1 R132H perturbs genome-wide expression patterns by increasing production of the oncometabolite D-2-hydroxyglutarate^65^, which competes with endogenous a-ketoglutarate to alter the activities of enzymes that are required to maintain normal DNA methylation^66^. Our findings support previous studies indicating that altered DNA methylation patterns can persist and perpetuate malignant phenotypes despite loss of the mutated protein that caused them^42–44^. This example is illustrative because it highlights the limitations of currentl gene panels for cancer diagnostics, which provide binary calls for the presence or absence of common oncogenic mutations. In this case, such panels would indicate the presence of IDH1 R132H and recommend treatment that targets this mutation^67^. However, with more precise knowledge of this tumor’s evolutionary history, we can see that such treatment will fail for one-fifth of malignant cells, since the mutated IDH1 protein is no longer there. In this case, it is these cells that will likely form the basis for therapeutic resistance.

There are several important methodological implications and limitations of our study. First, each tumor specimen analyzed in this study represents a fraction of overall tumor volume; future efforts will analyze multiple, geographically distinct tumor subsamples to evaluate the consistency of clonal architecture. Second, MOMA requires many sections to detect meaningful correlations (e.g., 25 sections provide ∼85% power to detect moderate correlations [|r| > 0.5, P < .05])^68^. Third, DNA and RNA must be co-isolated from each section (i.e., from the same population of cells). Fourth, deep sequencing is required to establish high-confidence VAFs for SNVs, which are in turn required to estimate clonal frequencies. Fifth, limited variability in clonal frequencies may impact the ability to detect corresponding molecular signatures. Sixth, some types of mutations are not yet captured by our approach (e.g., noncoding SNVs, rearrangements, chromothripsis, etc.). And seventh, collinearity in the abundance of malignant and / or nonmalignant cell types may produce spurious correlations (which can be mitigated by differential coexpression analysis with normal tissue, as done for case one, or sectioning in multiple planes, as done for case two). For this reason, we also recommend validating transcriptional profiles of malignant clones using one or more orthogonal techniques. We found that multiscale integration of bulk sections and single nuclei allowed us to leverage the complementary strengths of each sampling strategy. Specifically, bulk sections facilitate multiomic integration while yielding robust molecular signatures driven by millions of cells, while single nuclei enable precise validation of predictions made from bulk data. However, the success of this approach depends on accurate classification of malignant nuclei. In our study, we found that popular algorithms for identifying cancer cells from inferred CNVs from gene expression data^25,49,50^ were only ∼60% accurate. Therefore, MOMA will benefit from improved algorithms for inferring malignancy and / or scalable methods for profiling gene expression and malignant cell genotypes in parallel.

## Methods

### Pseudobulk analysis of scRNA-seq data

Single-cell RNA-sequencing (scRNA-seq) data from Venteicher et al.^24^ comprising 6243 cells from 10 IDH-mutant adult astrocytomas were downloaded from Gene Expression Omnibus (https://www.ncbi.nlm.nih.gov/geo/; accession ID = GSE89567). To generate a pseudobulk gene expression matrix from these data, 10% of all cells were randomly sampled and expression levels were summed for each gene from all sampled cells (this process was repeated 100x to generate a matrix with 100 pseudobulk samples). Using cell-class labels provided by the authors, the identities of all cells comprising each pseudobulk sample were tracked. Genome-wide differential expression analysis was performed by comparing all sampled malignant cells to all sampled nonmalignant cells using a two-sided t-test. In parallel, genome-wide gene coexpression analysis was performed as described^31^. Briefly, genome-wide biweight midcorrelations (bicor) were calculated using the WGCNA R package^69^ and all genes were clustered using the flashClust^70^ implementation of hierarchical clustering with complete linkage and 1 – bicor as a distance measure. The resulting dendrogram was cut at a static height of 0.277, corresponding to the top 1% of bicor values. All clusters consisting of at least 10 genes were identified and summarized by their module eigengene^52^ (i.e., the first principal component obtained by singular value decomposition) using the moduleEigengenes function of the WGCNA R package^69^. Highly similar modules were merged if the Pearson correlation of their module eigengenes was > 0.85. This procedure was performed iteratively such that the pair of modules with the highest correlation > 0.85 was merged, followed by recalculation of all module eigengenes, followed by recalculation of all correlations, until no pairs of modules exceeded the threshold. The pseudobulk gene coexpression module most strongly associated with malignant cells was identified by maximizing the correlation between the module eigengene and the actual fraction of sampled malignant cells in each pseudobulk sample. Genome-wide Pearson correlations to this module eigengene (*k*_ME_ values)^52^ were then calculated and compared to the results of single-cell differential expression analysis (t-values).

### Sample acquisition

The tumor specimen from case one (WHO grade II primary astrocytoma, IDH-mutant) was obtained from a 40 y.o. female patient following surgical resection at the University of California, San Francisco (UCSF), along with the patient’s blood (UCSF case ID: SF9495). The tumor specimen from case two (WHO grade II recurrent astrocytoma, IDH-mutant) was obtained from a 58 y.o. male patient following surgical resection at UCSF, along with the patient’s blood (UCSF case ID: SF10711). Four postmortem control human brain samples from two brain regions (anterior cingulate cortex [ACC] and entorhinal cortex [EC]) were also obtained from routine autopsies of two individuals (41 and 75 y.o. females) at UCSF. Control samples were examined by a neuropathologist (E.J.H.) and found to exhibit no evidence of brain disease. Tissue samples for nucleic acid isolation were immediately frozen on dry ice without fixation. For tumor histology, a smaller subsample was formalin-fixed and paraffin-embedded (FFPE) using standard procedures. All tumor samples were obtained with donor consent in accordance with protocols approved on behalf of the UCSF Brain Tumor Center Tissue Core.

### Serial sectioning

Tissue cryosectioning was performed on a Leica CM3050S cryostat at −20°C. Each sample was oversectioned to account for the possibility of low RNA quality or quantity from some cryosections; after excluding these (see below), most, but not all, analyzed sections were adjacent to one another. For the first case, 81 sections were cut and utilized as shown in **Fig. 2f**. For each of the four control samples, ∼120 sections were cut and 94 were utilized for gene expression profiling. For the second case, 140 sections were cut and utilized as shown in **Fig. 4f**. In addition, the plane of sectioning for the second case was rotated 90 degrees at the halfway point to provide additional spatial variation (**Fig. 4f**). These sectioning strategies resulted in 73% power to detect weak correlations (|r| > 0.3, P < .05) for case one and 83% power for case two^68^. To control for differences in the cross-sectional area of each tissue sample, section thickness was varied as needed to ensure sufficient and comparable amounts of nucleic acids could be extracted from sections for multiomic analysis. Quality control and usage information for all sections can be found in **Table S1** (case one), **Table S12** (control samples), and **Table S15** (case two). Frozen sections were collected in RNase-free 1.7 ml tubes (Denville Scientific Inc, South Plainfield, NJ) and stored at −80°C.

### Nucleic acid isolation and quality control

Tissue cryosections were thawed on ice and homogenized by pipette in QIAzol (Qiagen Inc., Valencia, CA). For control samples, RNA was extracted from each section with the miRNeasy mini kit (Qiagen Inc., Valencia, CA). For tumor samples, DNA and RNA were isolated simultaneously from each section with the AllPrep DNA / RNA / miRNA kit (Qiagen Inc., Valencia, CA). All nucleic acid isolation from tissue sections was performed using a QIAcube automated sample preparation system according to the manufacturer’s instructions (Qiagen Inc., Valencia, CA). Sections were processed in random batches of 12 on the QIAcube to avoid confounding section number with potential technical sources of variation associated with nucleic acid isolation.

Frozen blood was thawed and resuspended in red blood cell lysis solution (Qiagen Inc., Valencia, CA). White blood cells were removed by centrifugation at 2000g for 5 mins and repeated until white blood cells were depleted. Remaining red blood cells were resuspended in extraction buffer (50 mM Tris [pH8.0], 1 mM EDTA [pH8.0], 0.5% SDS and 1 mg / ml Proteinase K [Roche, Nutley, NJ]) and incubated overnight at 55°C. The extracted DNA was RNAse treated (40 μg / ml) (Roche, Nutley, NJ) for 1 h at 37°C before being phenol chloroform extracted and ethanol precipitated. The resulting DNA was resuspended in TE buffer (10 mM 460 Tris, 1 mM EDTA [pH7.6]).

RNA and DNA were analyzed using a Nanodrop 1000 spectrophotometer (Thermo Scientific Inc., Waltham, MA) to quantify concentrations, OD 260 / 280 ratios, and OD 260 / 230 ratios. Further validation of RNA and DNA concentrations was performed using the Qubit RNA HS kit and Qubit dsDNA HS kit on the Qubit 2.0 Fluorometer (Life Technologies Inc., Carlsbad, CA). RNA integrity (RIN) was assessed using an Agilent 2100 Bioanalyzer (Agilent Technologies Inc., Santa Clara, CA). Sections for which RIN ≥ 5 (case one median = 7.6, case two median = 8.3), OD 260 / 280 ratio ≥ 1.80 (case one median = 2.03, case two median = 1.94), and concentration by Nanodrop ≥ 9 ng / μl (case one median = 25.4 ng / μl, case two median = 9.25 ng / μl) were selected.

### Whole exome sequencing (WES) and data preprocessing

WES was performed at the UCSF Institute for Human Genetics genomics core facility (San Francisco, CA). Exome libraries were prepared from 1 μg of genomic DNA from each analyzed section using the Nimblegen EZ Exome kit V3 (Roche, Nutley, NJ). Paired-end 100 bp sequencing was performed on a HiSeq2500 sequencer (Illumina Inc., San Diego, CA). The analysis of WES data was performed as previously described^71^. Briefly, paired-end sequences were aligned to the human genome (University of California, Santa Cruz build hg19) using the Burrows-Wheeler Aligner (BWA)^72^. Uniquely aligned reads were further processed to achieve deduplication, base quality recalibration, and multiple sequence-realignment with the Picard suite^73^ and Broad Institute Genome Analysis ToolKit (GATK)^74^. After processing, a mean coverage of 131-151x and 104-122x was achieved for case one and case two, respectively.

### Single-nucleotide variant (SNV) and small insertion / deletion (indel) calling workflow

SNVs were identified using MuTect^75^ and indels were identified with Pindel^76^ using default settings. SNVs were further filtered to only retain variants with frequency > 0.10 in at least one tumor section and < 6 variant reads in the patient’s blood. Indels were filtered to only retain variants with > 5 variant reads in a given tumor section and < 13 total reads in the patient’s blood. If multiple indels were detected at the same genomic location, only the indel with the most supporting reads was retained. Identified mutations were annotated for their mutational context using ANNOVAR^77^ and were also cross-referenced with dbSNP^78^ (Build ID: 132) and the 1000 Genomes^79^ (Phase 1). SNV and indel events were converted to hg38 coordinates and assigned HGVS compliant names using Ensembl’s Variant Effect Predictor^80^.

### Droplet Digital PCR (ddPCR)

Variant allele frequencies (VAFs) of the IDH1 R132H mutation were determined in 69 tumor sections from case one and the patient’s blood using the PrimePCR IDH1 R132H mutant assay and the QX100 Droplet Digital PCR system (Bio-Rad Inc., Hercules, CA). An initial serial dilution of a positive control was performed to optimize the input concentration of genomic DNA from each section and to assess the reliability of the assay. Duplicate reactions were performed to quantify the reproducibility of the assay (**Fig. S1b**). Data were analyzed and 95% Poisson confidence intervals were calculated using QuantaSoft software (Bio-Rad Inc., Hercules, CA).

### Amplicon sequencing (amp-seq) and data preprocessing

Groups of mutations with similar allele frequency distributions in WES data were identified by hierarchical clustering. Biweight mid-correlations (bicor) were used to estimate the proximities of somatic mutations and 1-bicor was used as a dissimilarity measure. A subset of representative mutations from distinct clusters was validated by Sanger sequencing and deep sequencing of PCR amplicons (amp-seq) derived from tumor sections and the patient’s blood. Primers were designed using Primer-BLAST^81^ to yield an amplicon of around 500 bp (case one) or 100 bp (case two) with the mutation located within the center of the amplicon (**Tables S3 and S17**). Amplicons were generated for 42 mutations in case one (n = 69 sections; **Table S3**) and 75 mutations in case two (n = 85 sections; **Table S17**). For case one, the mutation-containing region was amplified by PCR using the FastStart high-fidelity PCR system (Roche, Nutley, NJ) or the GC-Rich PCR system (Roche, Nutley, NJ) as instructed by the manufacturer using specific annealing temperatures (**Table S3**). The resulting amplicons were purified using the NucleoSpin gel and PCR cleanup kit following the manufacturer’s instructions (Macherey-Nagel Inc., Bethlehem, PA) and submitted for Sanger sequencing with the same primers used to generate the amplicons. For case two, 50ng of gDNA was used as template per sample in each reaction and 35 cycles of PCR amplification were performed with KAPA HiFi HotStart Ready Mix (2x, KAPA Biosystems, Wilmington, MA). Multiplexed PCR reactions were purified using a 2X volume ratio of KAPA pure SPRI beads (KAPA Biosystems, Wilmington, MA). Purified PCR reactions were quantified using the Qubit dsDNA HS kit and Qubit 2.0 fluorometer. For both cases, the concentration of each amplicon was adjusted to 0.2 ng/μl. Barcoded libraries for each section were generated using the Nextera XT DNA Kit (Illumina Inc., San Diego, CA). After library preparation the barcoded libraries were pooled using bead-based normalization supplied with the Nextera XT kit. The pooled libraries were sequenced with paired-end 250 bp reads in a single flow cell on an Illumina MiSeq (Illumina Inc., San Diego, CA) in case one and an Illumina HiSeq 4000 in case two. In case one, libraries were sequenced in two runs, whereas all amplicons were sequenced in the same run for case two. Sequence reads were demultiplexed and basecalled using “bcl2fastq” (Illumina Inc., San Diego, CA). FASTQ files were aligned to a custom genome (based on the amplicon sequences) using BWA-MEM^82^. The SAMtools suite^83^ was used to create and index BAM files and create pileup files based on reads with a base quality score > 30. Read counts supporting the reference or variant within each amplicon were determined using the read counts function from VarScan 2^84^ and these counts were used to calculate VAFs.

### Downsampling analysis of amp-seq data

Amplicon reads originating from the reference or alternative alleles for *IDH1* or *TP53* were randomly downsampled to various coverage levels (n = 1000 random downsamples per coverage level) for each section to quantify the effect of reduced coverage on VAF estimates. VAFs were recalculated for each downsampled coverage level and compared to full coverage VAF estimates over all sections using Pearson’s correlation or root-mean-square error (RMSE), as illustrated in **Fig. S1c-d** (case one) and **Fig. S4a-b** (case two).

### Hierarchical clustering of variant allele frequencies (VAFs)

Groups of mutations with similar VAF patterns were identified by hierarchical clustering over all tumor sections. VAFs were clustered with Ward’s D method and 1 – Pearson’s correlation as a dissimilarity measure. The number of clusters was determined from the consensus of elbow^85^ and silhouette plot^86^ methods, using the cluster package in R^87^.

### DNA methylation data production and preprocessing

The sample order of genomic DNA from serial sections of case one was randomized to avoid confounding section number with potential sources of technical variation. DNA was concentrated with Genomic DNA Clean & Concentrator 10TM columns (Zymo Research, Irvine, CA) in batches of 12 samples, resulting in approximately two-fold concentration (median concentration after processing: 45ng / μl). The sample order was randomized again and concentrated DNA was shipped on dry ice to the University of California, Los Angeles (UCLA) Neurogenomics Core facility (Los Angeles, CA) for analysis using Illumina 450K microarrays (Illumina Inc., San Diego, CA).

Raw idat files were processed using the ChAMP R package^88^. Initial probe filtering was performed using the load.champ R function^89–91^. Probes with detection P-value > 0.01 (11,799 probes) or beadcount < 3 in at least 5% of samples were removed (n = 760), leaving 461,797 probes for analysis. The Illumina 450K microarrays contain two different assay types (Infinium I and Infinium II). Each assay has different sensitivity and dynamic range, which means that joint normalization leads to type II bias due to the lower sensitivity of the Infinium II assay^92^. We therefore performed beta-mixture quantile normalization (BMIQ) using the “champ.norm” function from ChAMP, which accounts for the different assay types^93^.

Additional preprocessing of the methylation data was performed with the SampleNetwork R function^94^, which identifies outlying samples, performs data normalization, and corrects for technical batch effects. The standardized sample network connectivity (Z.K) criterion was used to exclude one outlying sample (section #69, whose DNA concentration was substantially lower than other sections), leaving 68 sections. No batch effects associated with ArrayID or ArrayPosition were observed.

### Gene expression data production and preprocessing

Total RNA from case one (n = 69 sections) was shipped on dry ice to the UCLA Neurogenomics Core facility (Los Angeles, CA) for analysis using Illumina HT-12 v4 human microarrays (Illumina Inc., San Diego, CA). The order of the sections was randomized prior to shipment to avoid confounding potential technical artifacts with potential biological gradients of gene expression. Two control samples from the same pool of total human brain RNA (Ambion FirstChoice human brain reference RNA Cat#AM6050, Life Technologies Inc., Carlsbad, CA) were included with each of the five datasets. For each of the five datasets (case one and four control samples), all microarray samples (n = 72 – 96 / dataset) were processed in the same batch for amplification, labeling, and hybridization. Amplification was performed using the Ambion TotalPrep RNA amplification kit (Life Technologies Inc., Carlsbad, CA). Raw bead-level data were minimally processed by the UCLA Neurogenomics Core facility (no normalization or background correction) using BeadStudio software (Illumina Inc., San Diego, CA).

For each dataset the minimally processed expression data were further preprocessed using the SampleNetwork R function^94^. Using the standardized sample network connectivity (Z.K) criterion^94^, the following numbers of outliers were removed from each dataset: ACC1 (n = 2), ACC2 (n = 11), EC1 (n = 0), EC2 (n = 2), and case one (n = 1). Exclusion of outliers resulted in the following numbers of remaining sections in each dataset: ACC1 (n = 92), ACC2 (n = 83), EC1 (n = 94), EC2 (n = 92), and case one (n = 69). After removing outliers each dataset was quantile normalized^95^ and technical batch effects were assessed^94^. Significant batch effects (P < .05 after Bonferroni correction for univariate ANOVA) were corrected using the ComBat R function^96^ with no covariates as follows: ACC1 = ArrayID, ACC2 = ArrayID, EC1 = ArrayID and ArrayPosition, EC2 = QCBatch and ArrayID. No batch effects were observed for case one. Multiple technical batch effects were corrected sequentially. Analysis was restricted to 30,425 probes that were re-annotated^97^ as having either “perfect” (n = 29,272) or “good” (up to two mismatches; n = 1,153) sequence alignment to their target transcripts. Probes were further collapsed to unique genes (n = 20,019) by retaining one probe per gene with the highest mean expression over all sections.

For case two, RNA-sequencing was used to profile gene expression for all sections (n = 96). Full-length RNA was made into libraries using the KAPA stranded mRNA library prep kit (Roche, Nutley, NJ) following the manufacturer’s instructions, with a mean insert size of 300 bp. One ng of library (composed of library and ERCC spike-in controls, Life Technologies Inc., Carlsbad, CA) was added as input, and all libraries were normalized according to the manufacturer’s instructions. During this process samples were randomized in both section order and plane to avoid conflating biological and technical covariates. Sequencing was performed on eight lanes of a HiSeq4000 at the Center for Advanced Technology (CAT) at UCSF with single-end 50 bp sequencing using dual-index barcoding.

Reads were assessed with FastQC to ensure the quality of sequencing data by verifying high base quality scores, lack of GC bias, narrow distribution of sequencing lengths, and low levels of sequence duplication or adapter sequences^98^. Next, reads were subjected to adapter trimming using Cutadapt^99^ with minimum length = 20 and a quality cutoff of 20. Reads were subsequently aligned using default settings with the Bowtie2 program^100^ to the Genome Reference Consortium Human Build 37^101^. Finally, an expression matrix was generated using the FeatureCounts program with UCSC’s library of genomic features^102^ (n = 23,900 features). Genes with zero variance were removed (n = 30). Data were normalized with the RUVg package, regressing out 10 factors derived from principal component analysis of the ERCC spike-in control expression matrix^103^. The number of factors was determined empirically by evaluating relative log-expression (RLE) plots and gene-gene correlation distributions. Finally, the SampleNetwork R function^94^ was used to identify and remove six outlier sections based on the standardized sample connectivity criterion (Z.K).

### Copy number analysis by qPCR

The copy numbers for *TP53* and *ACCS* in case one were determined by SYBR Green-based qPCR. Primers were designed using Primer-BLAST^81^ and positioned immediately adjacent to but not including the SNV (ACCS F: TCTCTATGGCAACATCCGGC, R: CAGCCATGCAGCAACAGAAG; RPPH1 F: CGGAGGGAAGCTCATCAGTG, R: CCGTTCTCTGGGAACTCACC, TERT F: CTCGGATCATGCTGAGGACC, R: TTGTGCAATTCTGTGCCAGC, TP53 F: CAGTCACAGCACATGACGGA, R: GGGCCAGACCTAAGAGCAAT). qPCR was performed on genomic DNA from all 69 tumor sections and the patient’s blood using the LightCycler 480 SYBR Green I master mix and LightCycler 480 qPCR machine according to the manufacturer’s recommendations (Roche, Nutley, NJ). Measurements were triplicated and data were analyzed using the standard curve method. Copy numbers were determined for *TP53* and *ACCS* and two control genes on different chromosomes: ribonuclease P RNA component H1 (*RPPH1*) and telomerase reverse transcriptase (*TERT*) (data not shown). Relative copy number was determined by dividing the mean copy number of *TP53* and *ACCS* by the mean copy number of each reference gene separately to get a ratio and multiplying the ratio by two to obtain the diploid chromosome number. The relative copy number normalized to one of the reference genes (*RPPH1*) is shown in **Fig. S1e**.

### Copy number variation (CNV) calling (bulk data)

CNVs were quantified using multiple technologies and algorithms to generate reliable estimates. Although WES remains the gold-standard method for calling CNVs, DNA methylation and RNA-seq data provide cost-effective options that can be triangulated with sparse WES data to reduce false positives. Unless otherwise noted, default parameters were used. For case one we used the champ.CNA function, included with the ChAMP R package^39^, to call CNVs from DNA methylation data. For both cases, we called CNVs from exome data using FACETS^38^ with critical values of 25 (case one) and 450 (case two). Finally, we used CNVkit with circular binary segmentation to call CNVs from bulk RNA-seq data^45,104,105^.

### Generation of clonal trees with corresponding frequencies

CNVs were filtered to ensure that they were called in exome data and either DNA methylation data (case one) or RNA-seq data (case two) and covered more than 10% of a chromosomal arm. CNV coordinates were defined based on the intersection of ranges from both methods (**Tables S5** and **S19**). Using the frequencies of CNV / SNV mutations and tumor purity estimated from the *TP53* locus as input to PyClone^35^, we determined cluster membership for SNP and CNV events. We then used the PyClone output as the input to the CITUP algorithm^36^ to generate the most likely clonal tree (i.e., the tree with the minimum objective value) and derive clonal frequencies. In cases where there was an approximate tie between objective values, the tree was manually chosen based on biologically plausible principles. To visualize results we used the data.tree^106^ and DiagrammeR^107^ packages in R.

### Gene coexpression network analysis

Genome-wide biweight midcorrelations (bicor) were calculated using the WGCNA R package^69^ for case one (n = 20,019 genes) and case two (n = 23,870 genes). All genes were clustered using the flashClust^70^ implementation of hierarchical clustering with complete linkage and 1 – bicor as a distance measure. Each resulting dendrogram was cut at a static height (0.875 for case one and 0.562 for case two) corresponding to the top 30% and 20% of values of the correlation matrix for case one and case two, respectively. All clusters consisting of at least 15 members for case one or five members for case two were identified and summarized by their module eigengene^52^ (i.e. the first principal component obtained by singular value decomposition) using the moduleEigengenes function of the WGCNA R package^69^. Highly similar modules were merged if the Pearson correlation of their module eigengenes was > 0.80. This procedure was performed iteratively such that the pair of modules with the highest correlation > 0.80 was merged, followed by recalculation of all module eigengenes, followed by recalculation of all correlations, until no pairs of modules exceeded the threshold (case one: **Table S7**; case two: **Table S22**).

### Module enrichment analysis

The WGCNA measure of module membership, *k*_ME_, was calculated for all genes with respect to each module. *k*_ME_ is defined as the Pearson correlation between the expression pattern of a gene and a module eigengene and therefore quantifies the extent to which a gene conforms to the characteristic expression pattern of a module^52^ (case one: **Table S8**; case two: **Table S23**). For enrichment analyses, module definitions were expanded to include all genes with significant *k*_ME_ values, with significance adjusted for multiple comparisons by correcting for the false-discovery rate^108^. If a gene was significantly correlated with more than one module, it was assigned to the module for which it had the highest *k*_ME_ value. Enrichment analysis was performed for all modules using a one-sided Fisher’s exact test as implemented by the fisher.test R function.

### Lasso modeling of gene expression

The machine learning variable-selection method lasso (least absolute shrinkage and selection operator) and group lasso were performed using the R package Seagull^109–111^. Modeling was performed for each case with gene expression patterns as dependent variables and clonal frequency vectors as independent variables. For case one, clone 2 was excluded from modeling due to its low frequency and clone 6 was excluded since it was defined by a single CNV. Because clone 1 corresponds to the tumor purity vector, which represents the major vector of variation in this dataset, many genes experience inflated correlations to clone 1. To counteract this effect group lasso was performed. The truncal clone (clone 1) was placed in its own group and all remaining clones to be modeled were placed in a separate group. This procedure improved modeling performance for case one (**Fig. S2f-g**) but not case two (**Fig. S5f-g**), which may reflect the greater variance in tumor purity for case one. As such, modeling results for case two presented in the manuscript derive from the regular lasso model. For each gene, models were bootstrapped (n = 100) to address collinearity among clonal frequency vectors^112^ (as shown in **Fig. S2h** and **Fig. S5h**). We also generated empirical null distributions for model performance by permuting each gene’s expression profile prior to bootstrapping (n = 100).

When performing group-lasso modeling, only models with one surviving clonal frequency vector (not including the truncal clone) were considered. When performing lasso modeling, only models with one surviving clonal frequency vector were considered. To quantify model stability, we calculated the number of times out of 100 bootstraps that the most frequent surviving independent variable was the sole surviving variable. This stability metric was calculated for all gene models, including the permuted models. From the resulting distributions of stability values, a 5% FDR threshold was determined. For case one, the stability value of 73 represents the point beyond which 5% or fewer of the models were permuted models. Similarly, for case two the 5% FDR threshold for the stability metric was 45. Gene set enrichment analysis was performed via a one-sided Fisher’s exact test for all genes with significant model stability for the same clonal frequency vector (**Tables S6** and **S20** for case one and case two, respectively), with genes separated by the sign of the coefficient for the independent variable (**Tables S10** and **S24** for case one and case two, respectively).

### Differential gene coexpression analysis

Using the WGCNA R package^69^, pairwise biweight midcorrelations (bicor) were calculated among all 30,425 high-quality probes over all sections (n = 69–94) in each of five datasets (case one + four normal human brain samples), generating five identically proportioned correlation matrices (30,425 × 30,425). These correlations were then scaled to lie between [0,1] using the strategy of Mason et al.^113^. To identify gene coexpression relationships that were present in tumor but absent or weaker in normal human brain, each scaled bicor matrix produced from normal human brain was subtracted^114^ from the scaled bicor matrix produced from case one, resulting in four “subtraction matrices’’, or SubMats. The consensus of the four SubMats was formed by taking the minimum value at each point in the four matrices using the parallel minimum (pmin) R function, and the resulting “Consensus SubMat’’ was used as input for gene coexpression analysis (**Fig. S3a**). By definition, gene coexpression modules identified with this strategy will consist of groups of genes with expression patterns that are highly correlated in the astrocytoma but not in any of the normal human brain samples (**Fig. S3a**).

Probes in the Consensus SubMat were clustered using the flashClust^70^ implementation of a hierarchical clustering procedure with complete linkage and 1 – Consensus SubMat as a distance measure. The resulting dendrogram was cut at a static height of ∼0.38, corresponding to the top 2% of values in the Consensus SubMat. All clusters consisting of at least 10 members were identified and summarized by their module eigengene^52^ using the moduleEigengenes function of the WGCNA R package^69^. Highly similar modules were merged if the Pearson correlation of their module eigengenes was > 0.85. This procedure was performed iteratively such that the pair of modules with the highest correlation > 0.85 was merged, followed by recalculation of all module eigengenes, followed by recalculation of all correlations, until no pairs of modules exceeded the threshold. The WGCNA^69^ measure of intramodular connectivity (*k*_ME_) was calculated for all probes (n = 47,202) with respect to each module by correlating each probe’s expression pattern across all 69 tumor sections with each module eigengene.

### Single-nucleus DNA-sequencing and analysis

Three sections from case two (sections 29 and 113 / 115, which were combined) were analyzed by MissionBio, Inc. (MissionBio, San Francisco, CA) using their Tapestri microfluidics platform for single-nucleus DNA amplicon sequencing^46^. Using an in-house protocol, 4,433 (section 29) and 3,736 (sections 113 / 115) nuclei were extracted and recovered for analysis with the Mission Bio AML panel, which includes primers flanking one *IDH1* and two *TP53* loci. In addition, chr17 chromosomal copy number changes and *TP53* zygosity were inferred from a germline heterozygous intronic mutation upstream of TP53 G245V that happened to fall within the targeting panel (NC_000017.11:g.7674797T>A). Sequencing was performed on a MiSeq (Illumina Inc., San Diego, CA), yielding an average of 6,801 (section 29) or 6,433 (sections 113 / 115) reads per nucleus, with alignment rates of ∼90%. Hierarchical clustering of nuclei for mutations of interest was performed separately for section 29 and sections 113 / 115 using complete linkage and Euclidean distance, with *k* = 4 chosen based on silhouette^86^ and elbow plots^85^. Genotype calls for the clusters were manually annotated as described in **Fig. S4f** and **Table S21**.

### Single-nucleus RNA-sequencing and analysis

#### Library prep and sequencing

Four sections (17, 53, 93, 117) from case two were used to generate single-nucleus RNA-seq (snRNA-seq) data. Our approach was adapted from TARGET-Seq^47^, a protocol utilizing dual-indexing of sample barcodes and unique molecular identifiers (UMIs) of captured transcripts. Briefly, for each section, lysis was performed by dounce homogenization with staining of nuclei by Hoescht3342 and subsequent flow-sorting into three 96-well plates per section. Each plate was randomized and subsequently processed individually and in random order. We used the SmartScribe kit (Takara Bio USA, San Diego, CA) for RT-PCR, followed by PCR with the SeqAmp PCR kit (Takara Bio USA, San Diego, CA). Unlike TARGET-Seq, the RT reaction was performed using only polyA primers (**Table S26**). ERCC spike-in control RNA was added to the wells according to manufacturer’s instructions to facilitate identification and correction of batch effects. Wells for each plate were pooled in equivolume proportions and an Agilent 2100 Bioanalyzer (Agilent Technologies Inc., Santa Clara, CA) was used to assess sample quality and cDNA concentrations were quantified using a Qubit 2.0 Fluorometer with the dsDNA-High Sensitivity kit (Life Technologies Inc., Carlsbad, CA), yielding mean cDNA concentration of 1ng/ul. Concentrations were normalized prior to tagmentation (Nextera Kit, Illumina Inc., San Diego, CA) and amplification of 3’ ends, as in TARGET-Seq^47^. Sequencing was performed using the 150-cycle high-throughput kit on an Illumina NextSeq550 at SeqMatic (Fremont, CA) with dual-indexed sequencing and read parameters as in TARGET-seq.

#### Data preprocessing

snRNA-seq raw reads were demultiplexed and basecalled using “bcl2fastq” (Illumina Inc., San Diego, CA). Barcodes were filtered using the “umi_tools” package^115^ whitelist function, with a Hamming distance of 2 and the density knee method to determine the number of true barcodes. 809 / 1152 nuclei (70.2%) passed this initial quality control step. Reads were assessed with FastQC to ensure the quality of sequencing data by verifying high base-quality scores, lack of GC bias, narrow distribution of sequencing lengths, and low levels of sequence duplication or adapter sequences^98^. Next, reads were subjected to adapter trimming using the Trimmomatic algorithm^116^ with a minimum length of 30, a minimum quality of 4 with a 15 bp sliding window, and otherwise default settings. A mean of 445,082 reads / nucleus was achieved at this stage. Reads were subsequently aligned using ENCODE RNA-seq settings (except for outFilterScoreMinOverLread, which was set to 0) with the STAR program^117^ to the Genome Reference Consortium Human Build 38^101^. Finally, an expression count matrix was generated using the FeatureCounts program^118^ with Gencode’s library of gene features (version 21)^119^, subset using the “gene” attribute (n = 60,708 features). Deduplication of UMIs was performed using a custom R script, resulting in a mean number of 206,638 unique reads / nucleus, a 46% deduplication rate. Features with counts less than one in more than 90% of cells were removed (n = 57,021 final features). Data were normalized with the RUVg package, regressing out 10 factors derived from PCA of the ERCC spike-in control expression matrix^103^. Normalized counts were further processed using the Sanity package^27^, with 1000 bins and a minimum and maximum variance of 0.001 and 1000, respectively. Internuclear distance was determined using the Sanity_distance function with a signal to noise parameter of 1 and inclusion of error bars.

#### snRNA-seq clustering and differential expression analysis

snRNA-seq data were hierarchically clustered using the hclust function in R with Ward’s method and the distance metric derived by Sanity^27^. This distance metric uses a Bayesian approach by giving less weight to gene expression estimates with large error bars when calculating cell distances. The optimal number of clusters (*k* = 12) was determined using elbow^85^ and silhouette plots^86^ with the cluster package in R^87^. Differential expression analysis (t.test) was performed between each cluster and all other clusters using Sanity-adjusted expression values for all genes. The resulting distributions of t-values were then compared for genes comprising the bulk coexpression modules most strongly associated with each malignant clone / nonmalignant cell type and all other genes (white and black distributions, respectively, in **Fig. S6a-j**; significance was evaluated with a one-sided Wilcoxon rank-sum test). Module genes were defined as those that were significantly correlated with corresponding bulk coexpression module eigengenes as determined by the FDR threshold^108^. If a gene was significantly correlated with more than one module eigengene, it was assigned to the module for which it had the highest *k*_ME_ value.

#### CNV calling

The snRNA-seq count matrix was used as input to CopyKat^49^. Nuclei snRNA-seq clusters determined to be non-malignant by snAmp-seq were used as normal control cells. “KS.cut” was set to 2, “ngene.chr” was set to 20, and Ensembl gene names were used. InferCNV^25^ was provided with a vector of nonmalignant cells (as previously determined) based on clustering and snAmp-seq in “subclusters” mode, with a cutoff parameter of 1, and denoising turned on, “ward.D” as clustering method, “qnorm” as subcluster partition method, and tumor subcluster p-value of 0.05. The Hidden Markov model was not used. The program CaSpER^50^ was run with the raw snRNA-seq count matrix as input and default settings, again using snRNA-seq clusters of nuclei determined to be malignant by snAmp-seq as negative controls. For each of these algorithms, the outputs were clustered based on Euclidean distance using Ward’s D method. Clusters with no CNV signal were labeled nonmalignant while all other clusters were presumed to represent malignant cells.

Sensitivity and specificity were calculated using the snAmp-seq data as ground truth. True positives (TP) were defined as the intersection of malignant calls by the CNV calling algorithm and the snAmp-seq data. True negatives (TN) were defined as the intersection of nonmalignant calls by the CNV calling algorithm and the snAmp-seq data. False negatives (FN) and false positives (FP) were similarly defined. Nuclei with insufficient data were excluded from the analysis. Sensitivity was defined as: TP / (TP + FN), while specificity was defined as: TN / (TN + FP). Accuracy was defined as (TP + TN) / (TP + FP + TN + FN).

#### UMAP and trajectory analysis

UMAP was performed for all nuclei (n = 809) with a starting seed of 15, 30 neighbors, a spread of 3, a minimum distance of 2, and 1 – Pearson correlation as a distance metric using the “uwot” R package^120^ after selecting the first 30 principal components of the Sanity-corrected expression matrix including all genes. UMAP was also performed separately for all cells associated with malignant clusters using the Sanity-corrected expression matrix. After selecting the first 15 principal components, the “uwot” package was used with a seed of 15, 20 neighbors, a spread of 3, a minimum distance of 2, and 1-Pearson correlation as the similarity metric. All other settings were left as defaults. Trajectory analysis was performed with the Slingshot R package^51^ on the UMAP plot. The “simple” distance method was used and all other parameters were left as their default values.

#### Gene set enrichment analysis

Enrichment analysis (one-sided Fisher’s exact test) was performed for each snRNA-seq cluster using genes that were differentially expressed in that cluster relative to all other clusters using a one-sided Wilcoxon rank-sum test. Resultant p-values were further FDR-corrected to q-values^108^. Gene sets used for enrichment analysis are listed in **Table S9.**

#### Amp-seq genotyping

Single-nucleus amplicon-seq (snAmp-seq) of single-nucleus cDNA was adapted from the TARGET-seq protocol^47^. Primers flanking the following mutations (marking the truncal clone) were designed with Primer3^121^: IDH1 R132H, TP53 G245V, and RUFY1 K218N (**Table S26**). To overcome lack of heterogeneity in sequencing, random spacers were added to the beginning (5’ end) with 0 - 5 nucleotides from the sequence CGTAC. Finally, a common sequence was added to the 5’ end of the primer for a second round of PCR (**Table S26**). We selected wells that passed QC for snRNA-seq analysis and processed each plate separately and in random order. Amplification of the first round of PCR was performed with the KAPA 2G Ready Mix (Roche Inc., Nutley, NJ) with the same PCR program as for TARGET-Seq^47^. The program “Barcrawl”^122^ was used to create custom dual-index barcodes for the amplification PCR. At this stage 10% of wells were checked using an Agilent 2100 Bioanalyzer (Agilent Inc., Santa Clara, California) to determine whether products of appropriate size were produced. All wells were quantified with a Qubit 2.0 Fluorometer using the dsDNA-High Sensitivity kit (Life Technologies Inc., Carlsbad, CA) and normalized prior to the next step. The second round of PCR used custom sequencing primers that were partially complementary to the previous sequences, with custom dual-index barcodes generated from BarCrawl^122^ and Illumina P5 / P7 sequences. Sequencing was performed using a 300 cycle Miseq v2 Nano kit on a MiSeq (Illumina Inc., San Diego, CA).

snAmp-seq data were demultiplexed and basecalled using “bcl2fastq” (Illumina Inc., San HDiego, CA). Reads were assessed with FastQC to ensure the quality of sequencing data by verifying high base quality scores, lack of GC bias, narrow distribution of sequencing lengths, and low levels of sequence duplication or adapter sequences^98^. Next, reads were subjected to adapter trimming using the Trimmomatic algorithm^116^ with a minimum length of 30, a minimum quality of 4 with a 15 bp sliding window, and otherwise default settings^116^. Reads were subsequently aligned with the STAR program to a custom version of the genome containing only the amplicons of interest. Default parameters were altered such that no multiple alignments or splicing events were allowed. The median number of reads per nucleus for each amplicon was (IDH1 R132H: 177; TP53 G245V: 246; RUFY1 K218N: 209). Read counts supporting the reference or variant allele within each amplicon were determined using the read counts function from VarScan 2^84^ and these counts were used to calculate variant frequencies. Nuclei were sorted into three categories: called nuclei (calls by VarScan 2 of two or more mutant or two or more wild-type [WT] calls of the three loci with either one or zero indeterminate calls), discrepant nuclei (two WT and one mutant call), and insufficient data nuclei (two or more loci in which VarScan 2 was unable to call a genotype). The breakdown for these categories is as follows: 75% called nuclei, 1% discrepant nuclei, and 24% insufficient data nuclei (**Table S27**).

### Inter-case analysis

Combined Pearson correlations to tumor purity for the 15,288 genes shared between case one and case two were determined by calculating the weighted average of the z-scores produced by Fisher’s transformation, dividing this value by the joint standard error, and applying the inverse Fisher transformation^31^. To define significant genes for enrichment analysis (**Fig. 9a-b**), a minimum absolute value for Pearson’s correlation of > 0.3 or < −0.3 was required in both cases along with an FDR-corrected q-value of < 0.05. Enrichment analysis was performed as described above, with gene sets listed in **Table S9**. Significant positively correlated genes were subjected to protein-protein interaction (PPI) analysis using the STRING database^53^. We used the STRINGdb^53,54^, network^123^, intergraph^124^, and ggnetwork^125^ packages to visualize the results of STRING PPI analysis. The “physical” network flavor and minimum score of 900 was utilized to guarantee that all depicted interactions were actual PPIs with experimental evidence. Clusters with more than five members were chosen from the set of interaction clusters generated from all genes that had positive correlations and passed the correlation cutoffs listed above. Enrichment analysis of PPI clusters was performed as described above, with gene sets listed in **Table S9**.

### Histology and immunostaining

Tumor tissue was fixed in 10% neutral-buffered formalin, processed, and embedded in paraffin. Tumor sections (5 μm) were prepared and stored at −20°C prior to use. Hematoxylin and eosin staining was performed using standard methods. As part of clinical evaluation, the proliferative index and TP53 mutation status were estimated based on review of immunostained slides for KI67 or TP53, respectively. Briefly, in regions with increased signal the percent of tumor cells staining was estimated based on review of ten 200x fields.

Anti-AKR1C3 was selected based on statistical considerations pursuant to bioinformatic analyses and after preliminary validation of efficacy in human tissue via The Human Protein Atlas^126^ (http://www.proteinatlas.org). Primary antibodies and conditions were IDH1 R132H (DIA-H09, Dianova, mouse clone H09, dilution 1:50); AKR1C3 (Catalog# AB84327, Abcam, rabbit polyclonal, dilution 1:600 for single immunohistochemistry and 1:1200 for dual immunofluorescence); and TP53 (1:25, Novocastra, catalog # P53-D07-L-CE-H). Heat antigen retrieval was performed in Tris-EDTA at pH8. Following antigen retrieval, sections for immunohistochemistry were treated with 3% methanol-hydrogen peroxide at 22°C for 16 min.

All immunostaining and multiplex immunostainings were performed using a Discovery XT autostainer or Benchmark XT (Ventana Medical Systems, Inc., USA). For signal detection, the Multimer HRP kit (Ventana Medical Systems, Inc., USA) followed by either DAB or fluorescent detection kits were used. Fluorophores with the least autofluorescence on FFPE tissue were selected to minimize false positives: Cyanine 5 (Cy5) (DISCOVERY CY5 Kit, Cat#760238, Roche Diagnostics Corporation, Indianapolis, USA) and rhodamine (DISCOVERY Rhodamine Kit, Cat#760233, Roche Diagnostics Corporation, Indianapolis, USA). Slides were then counterstained with DAPI (Sigma Aldrich, USA) at 5 μg/ml in PBS (Sigma Aldrich, USA) for 15 minutes, mounted with prolong Gold antifade mounting media reagent (Invitrogen, USA) and stored at −20°C prior to imaging. Positive and negative controls were included for each marker. Images of stained slides were acquired using either a light microscope (Olympus BX41 microscope using UC90 Cooled CCD 9 Megapixel camera) or Zeiss Cell Observer epifluorescence microscope equipped with an AxioCam 506M camera and an Excellitas X-Cite 120Q light source and processed with Photoshop CS6 (Adobe systems, San Jose, CA). Nonmalignant tissue analyzed in **Fig. 9** was obtained from a patient with epilepsy and corresponds to normal tissue adjacent to epileptic foci.

### Data analysis and figure production

Unless otherwise stated, all analyses were performed in the R computing environment (https://www.r-project.org). Figures were produced with the aid of the R packages ggplot2^127^, data.table^128^, RColorBrewer^129^, gridExtra^130^, ComplexHeatmap^131^, Circlize^132^, and ggsignif^133^.

### Data and code availability

All data are publicly available for download under NCBI Bioproject ID PRJNA953039. Code for processing data and producing figures featured in this manuscript is available on GitHub: https://github.com/oldham-lab/Deconstructing-Intratumoral-Heterogeneity-through-Multiomic-and-Multiscale-Analysis.../tree/main

## Supplementary Figure and Table Legends

**Figure S1.**
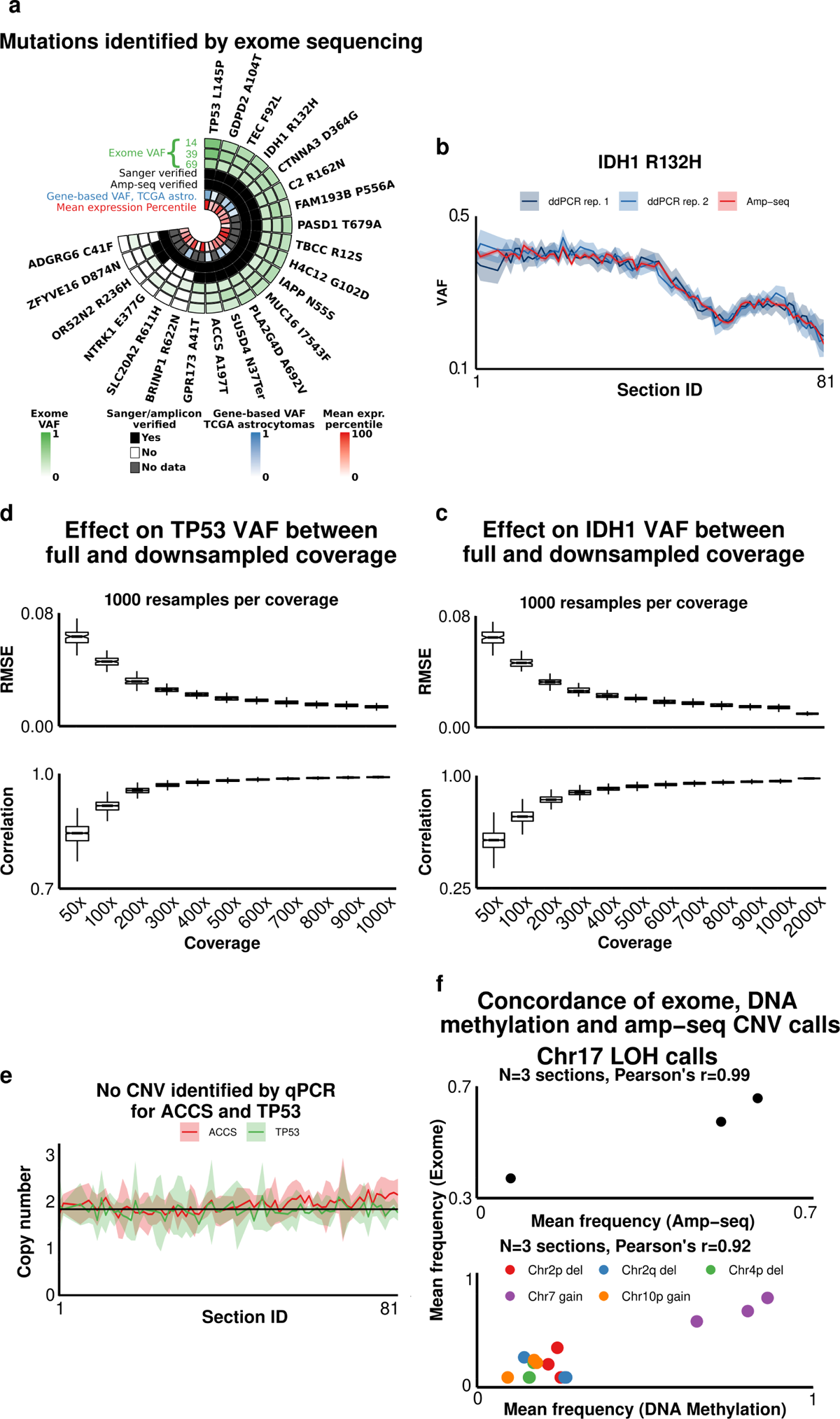
Mutation validation (case 1). **a)** Nonsynonymous mutations were identified by exome sequencing of tumor sections 14, 39, 69, and the patient’s blood. Green track: variant allele frequencies (VAF) for each mutation in each section. Black tracks: mutation validation by Sanger sequencing and amp-seq, which is more sensitive. Blue track: gene mutation frequencies in TCGA astrocytomas (n = 286). Red track: mean expression percentiles for each gene over all tumor sections. **b)** Amp-seq and droplet-digital PCR (ddPCR) yielded consistent estimates of IDH1 R132H variant frequencies (n = 69 tumor sections; rep. 1 and rep. 2 denote technical replicates using the same input DNA). Shaded areas represent two standard errors. **c-d)** Downsampling of amp-seq reads for IDH1 R132H **(c)** and TP53 L145P **(d)** was performed in each tumor section to achieve desired coverage levels (x-axis). For each downsampling (n = 1,000), the root mean square-error (RMSE; top) and Pearson’s correlation (bottom) was calculated with respect to the true VAF (calculated using all reads) over all sections (n = 69). **e)** Relative copy number was determined by SYBR Green qPCR for *TP53* and *ACCS* loci using genomic DNA from 69 tumor sections and blood. The mean of triplicate measurements, normalized to RNaseP (*RPPH1*) copy number, is shown. Shaded areas represent two standard errors. **f)** Top: Concordant estimates of chr17p loss-of-heterozygosity (LOH) in the same tumor sections (n = 3) were obtained from exome data by analyzing changes in B-allele frequencies and from amp-seq data by analyzing TP53 L145P VAF, which is equivalent to chr17p LOH frequency since both events are truncal. Bottom: Concordant estimates of CNV frequencies in the same tumor sections (n = 3) were obtained using FACETS^38^ and ChAMPS^39^ to analyze exome and DNA methylation data, respectively.

**Figure S2.**
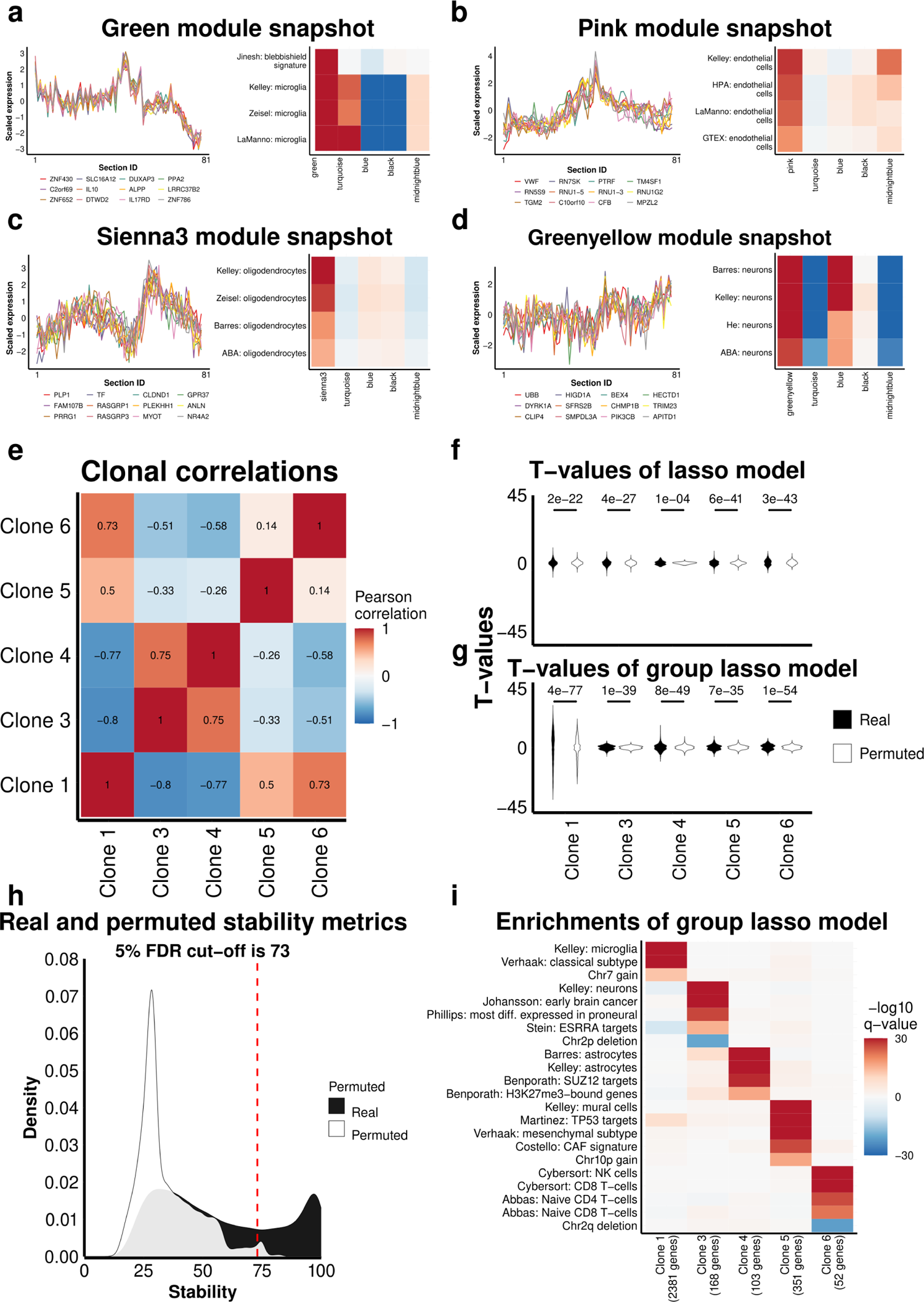
Linear modeling of gene expression using clonal frequencies reveals concordant gene-set enrichments with coexpression modules (case 1). **a-d)** Left: snapshots of additional gene coexpression modules enriched for markers of nonmalignant cell types (expression patterns for the top 12 genes ranked by *k*_ME_ are shown). Right: heatmaps of gene set enrichment results for each module. Modules included genes that were most specifically and significantly correlated (after FDR correction) to the module eigengene (ME), and enrichment was assessed with a one-sided Fisher’s exact test (followed by FDR correction; see panel **i** for legend). **e)** Correlation heatmap for the cumulative frequency vectors of identified clones. **f-g)** Lasso regression^109^ was used to model the expression of all genes (n = 20,018) as a function of clonal frequencies over all tumor sections (n = 69). Violin plots illustrate the distributions of t-values for all models where the indicated clone was the only explanatory variable that survived lasso selection. Permutations were performed by randomly scrambling clonal frequencies (n = 100) prior to lasso regression. Real and permuted clonal frequency vectors were bootstrapped (n = 100) to address collinearity. P-values denote the significance of the Anderson-Darling test, which evaluates whether two distributions are likely to be derived from the same distribution. **f)** Results of a standard lasso model. **g)** Results of a group lasso model where the truncal clone (equivalent to tumor purity) was placed in a separate group due to its strong effect on gene expression (Fig. 3c); note the general improvement in Anderson-Darling test P-values. **h)** Density plot showing the number of times (out of 100 bootstraps) that the same explanatory (clonal frequency vector) variable was retained by the group lasso regression model, or ‘stability’. Only group lasso models where retained explanatory variables included the truncal clone and up to one other clone were considered. The vertical line demarcates the point to the right of which only 5% of values belong to the permuted distribution, i.e. a 5% FDR rate. **i)** Enrichment analysis (one-sided Fisher’s exact test) of genes that were significantly (FDR < .05) and stably (FDR < .05) associated with each clone. Gene sets are described in **Table S9**. Heatmap depicts -log10 FDR-corrected p-values (q-values; shared legend for **a-d**) after comparing each gene set to all genes with stability > 73 for a given clone (one-sided Fisher’s exact test). Positive values represent enrichments for genes with significant positive correlations to the ME (**a-d**) or significant positive modeling coefficients (**i**), while negative values represent enrichments for genes with significant negative correlations to the ME (**a-d**) or significant negative modeling coefficients (**i**).

**Figure S3.**
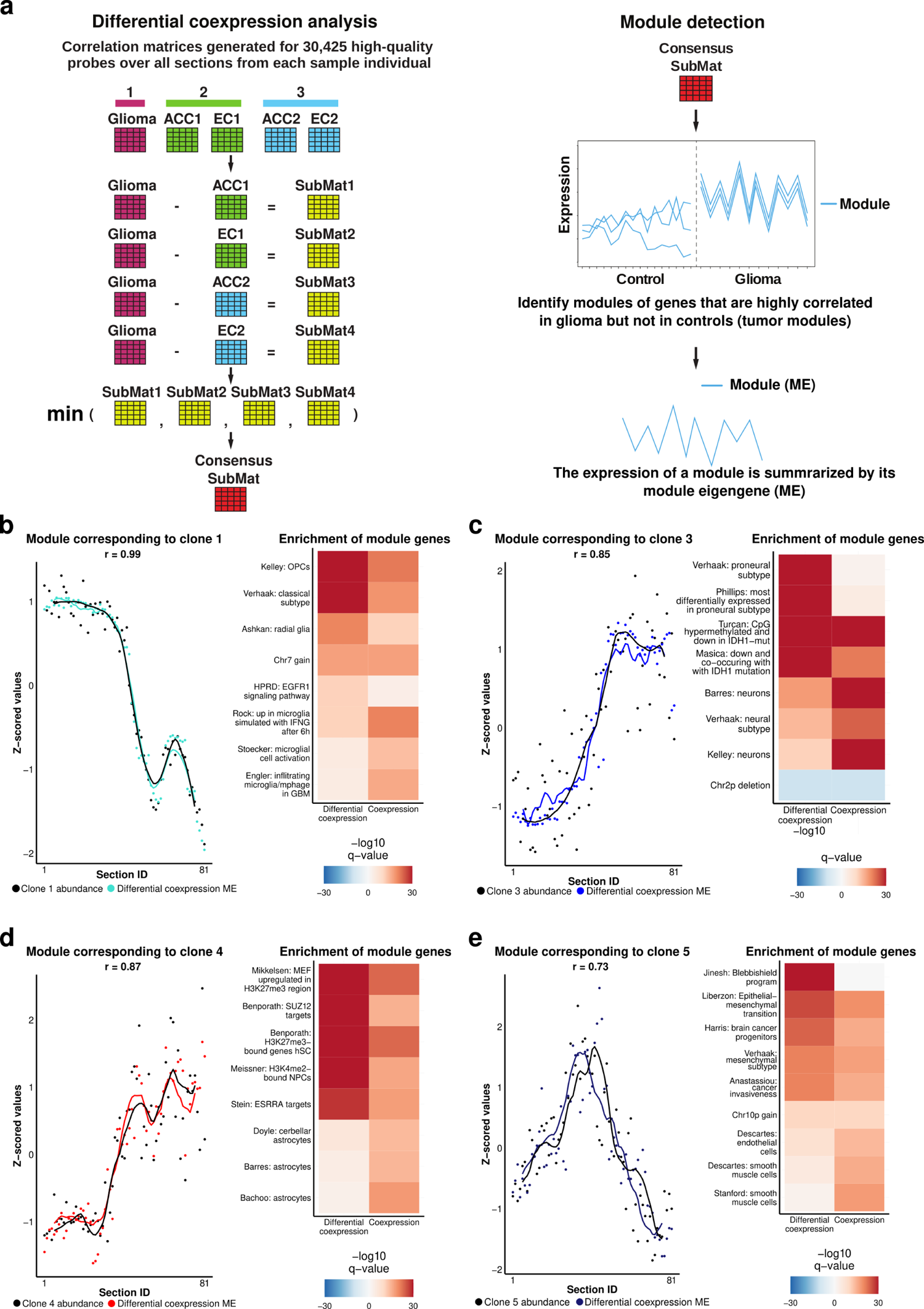
Differential coexpression analysis of glioma and normal human brain preserves gene coexpression modules associated with malignant clones (case 1). **a)** Genome-wide gene coexpression relationships were calculated for each of the five tissue specimens (one astrocytoma and four normal brain controls) over all tissue sections, resulting in five correlation matrices with the same dimensions. Unbiased differential coexpression analysis was performed as illustrated. ACC = anterior cingulate cortex; EC = entorhinal cortex. **b-e)** Left: differentially coexpressed module eigengenes (ME) with the strongest correlations to clonal abundance (defined cumulatively). Locally weighted smoothing (LOESS) lines are shown; correlation is based on data points. Right: enrichment analysis of differentially coexpressed module genes using published gene sets. FDR-corrected p-values (q-values) from one-sided Fisher’s exact tests are shown. Positive values represent enrichments of genes that were significantly positively correlated to the ME, while negative values represent enrichments of genes that were significantly negatively correlated to the ME. Gene sets representing chromosomal gains or losses include all genes within affected regions (as described in Fig. 2l and **Table S5**). See **Table S9** for descriptions and sources of featured gene sets.

**Figure S4.**
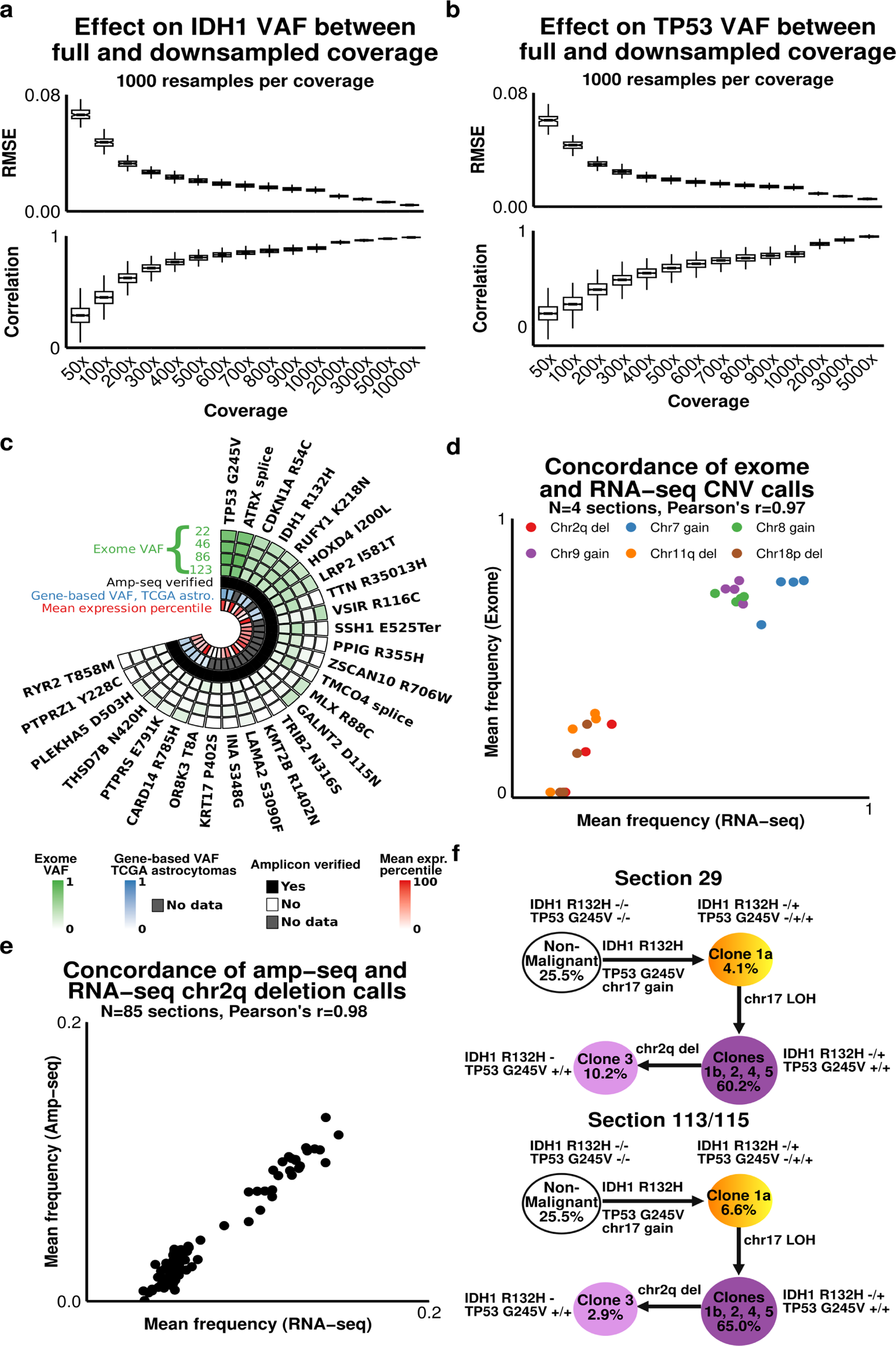
Mutation validation (case 2). **a-b)** Downsampling of amp-seq reads for IDH1 R132H (**a**) and TP53 G245V (**b**) was performed in each tumor section to achieve desired coverage levels (x-axis). For each downsampling (n = 1,000), the root mean square-error (RMSE; top) and Pearson’s correlation (bottom) was calculated with respect to the true VAF (calculated using all reads) over all sections (n = 85). **c)** Nonsynonymous mutations were identified by exome sequencing of tumor sections 22, 46, 85, 123, and the patient’s blood. Green track: variant allele frequencies (VAF) for each mutation in each section. Black track: mutations validation by amp-seq. Blue track: gene mutation frequencies in TCGA astrocytomas (n = 286). Red track: genome-wide mean expression percentiles over all sections (n = 90). **d)** Concordant estimates of CNV frequencies in the same tumor sections (n = 4) were obtained using FACETS^38^ and CNVkit^45^ to analyze exome and RNA-seq data, respectively. **e)** Concordant estimates of chromosome 2q deletion frequencies in the same tumor sections (n = 85) were obtained using amp-seq (Fig. 4m) and RNA-seq, which was analyzed by CNVkit. **f)** Clone phylogeny (with arbitrary branch lengths) derived from single-nucleus amp-seq (snAmp-seq) of mutations affecting the *IDH1* and *TP53* loci for section 29 (n = 4,433 nuclei) and sections 113/115 (n = 3,736 nuclei). Clone names are derived from Fig. 4o, and the percentages of nuclei assigned to each clone are shown.

**Figure S5.**
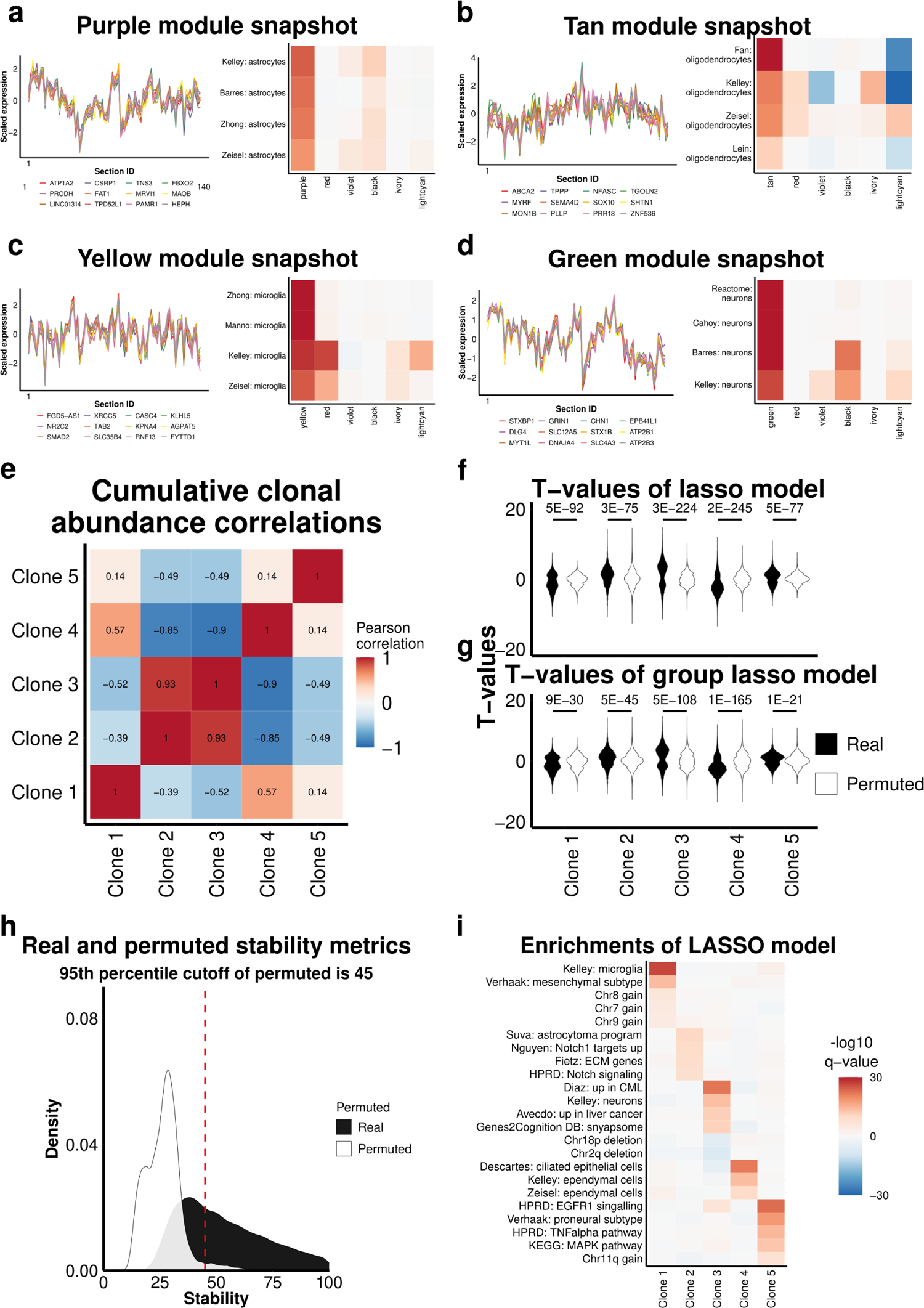
Linear modeling of gene expression using clonal frequencies reveals concordant gene-set enrichments with coexpression modules (case 2). **a-d)** Left: snapshots of additional gene coexpression modules enriched for markers of nonmalignant cell types (expression patterns for the top 12 genes ranked by *k*_ME_ are shown). Right: heatmaps of gene set enrichment results for each module. Modules included genes that were most specifically and significantly correlated (after FDR correction) to the module eigengene (ME), and enrichment was assessed with a one-sided Fisher’s exact test (followed by FDR correction; see panel **i** for legend). **e)** Correlation heatmap for the cumulative frequency vectors of identified clones. **f-g)** Lasso regression^109^ was used to model the expression of all genes (n = 20,246) as a function of clonal frequencies over all tumor sections (n = 85). Violin plots illustrate the distributions of t-values for all models where the indicated clone was the only explanatory variable that survived lasso selection. Permutations were performed by randomly scrambling clonal frequencies (n = 100) prior to lasso regression. Real and permuted clonal frequency vectors were bootstrapped (n = 100) to address collinearity. P-values denote the significance of the Anderson-Darling test, which evaluates whether two distributions are likely to be derived from the same distribution. **f)** Results of a standard lasso model. **g)** Results of a group lasso model where the truncal clone (equivalent to tumor purity) was placed in a separate group. Unlike case 1, the group lasso model did not outperform the standard lasso model. **h)** Density plot showing the number of times (out of 100 bootstraps) that the same explanatory (clonal frequency vector) was retained by the standard lasso regression model, or ‘stability’. The vertical line demarcates the point to the right of which only 5% of values belong to the permuted distribution, i.e. a 5% FDR rate. **i)** Heatmap of FDR-corrected p-values (q-values; shared legend for panels **a-d**) after comparing each gene set to all genes with stability > 45 for a given clone (one-sided Fisher’s exact test). Positive values represent enrichments of genes with significant positive correlations to the ME (**a-d**) or significant positive modeling coefficients (**i**), while negative values represent enrichments of genes with significant negative correlations to the ME (**a-d**) or significant negative modeling coefficients (**i**).

**Figure S6.**
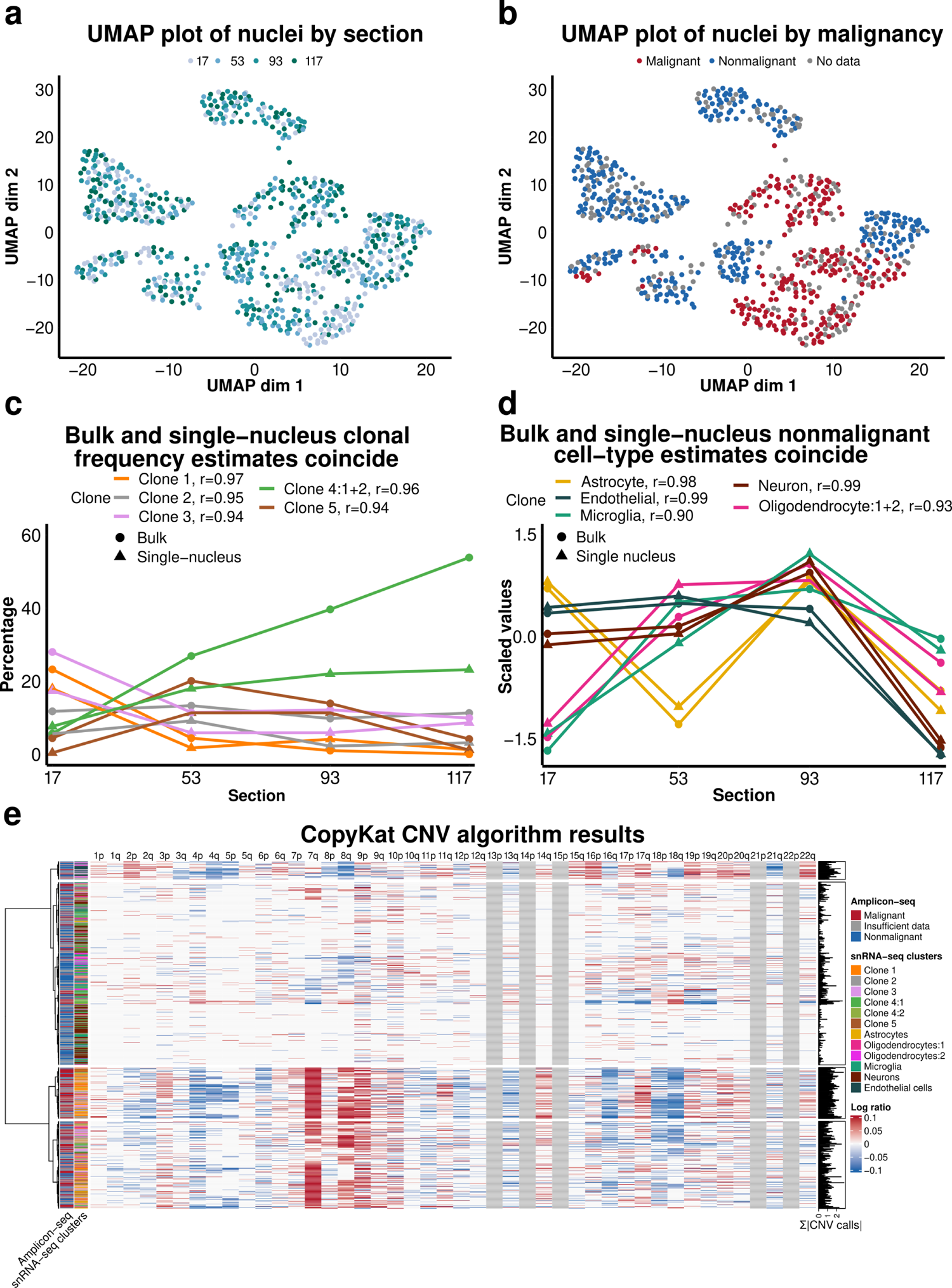
Single-nucleus RNA-seq analysis validates inferences from bulk data. **a)** UMAP plot of snRNA-seq data (n = 809 nuclei) with the tumor section IDs that served as the source for each nucleus superimposed. **b)** UMAP plot of snRNA-seq data with malignancy superimposed. Malignancy was determined by genotyping all nuclei via single-nucleus amplicon sequencing (snAmp-seq) of cDNA spanning mutations in the truncal clone. **c)** Frequencies of malignant clones in snRNA-seq data (n = 360 nuclei from four tumor sections) and bulk data (n = 16 tumor sections), with correlations in legend. **d)** Relative abundance of nonmalignant cell types in snRNA-seq data (n = 449 nuclei from four tumor sections) and bulk data (n = 16 tumor sections), with correlations in legend. Estimates were scaled and centered for comparability. Bulk estimates for (**c-d**) are derived from clonal abundance and module eigengene values featured in Fig. 4p and **Fig. S5a-d**, respectively, averaged across the four sections flanking each section analyzed by snRNA-seq (snRNA-seq section 17: bulk sections 14, 16, 18, 19; snRNA-seq section 53: bulk sections 50, 51, 54, 55; snRNA-seq section 93: bulk sections 91, 92, 94, 95; snRNA-seq section 117: bulk sections 114, 116, 118, 119). **e)** Log-ratio output of the CopyKat CNV algorithm^49^. Left: snAmp-seq malignancy assignments and snRNA-seq cluster assignments. Right: sum of the absolute value of CopyKat CNV calls (chromosomal arms in gray could not be called due to inadequate gene coverage).

**Figure S7.**
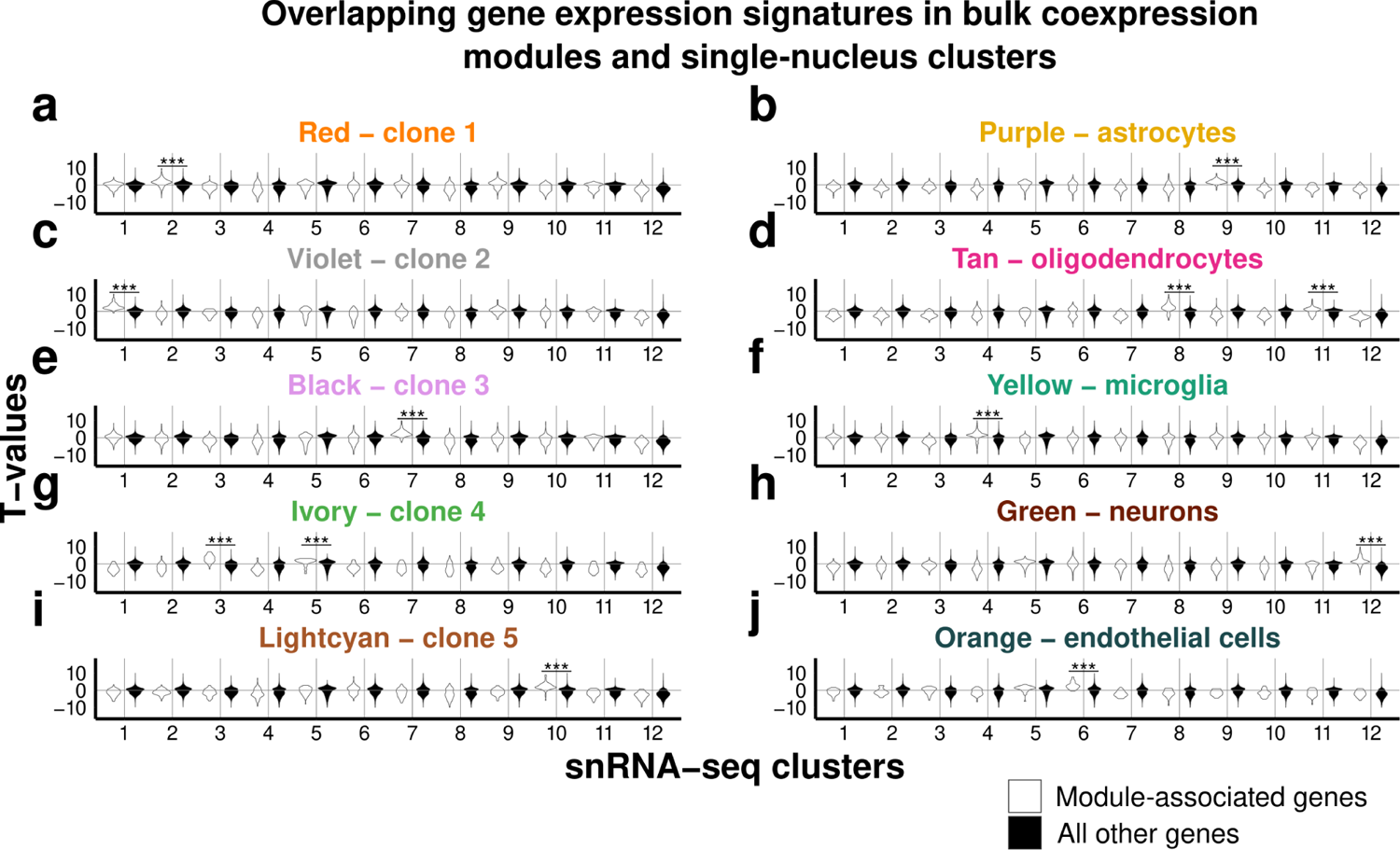
Bulk coexpression module genes map definitively onto single-nucleus clusters. **a-j)** Modules of coexpressed genes from bulk tumor sections (n = 90) that were most strongly associated with specific clones (Fig. 5d**-g**) or nonmalignant cell classes (**Fig. S5a-d**) were evaluated for differential expression in each snRNA-seq cluster vs. all other clusters (white distributions: t-test results for all module genes). Genes that were not associated with each module were evaluated in the same fashion (black distributions), and a one-sided Wilcoxon rank-sum test was used to determine whether module genes were significantly upregulated in a given snRNA-seq cluster relative to all other genes (*** = P < 1e-10).

**Figure S8.**
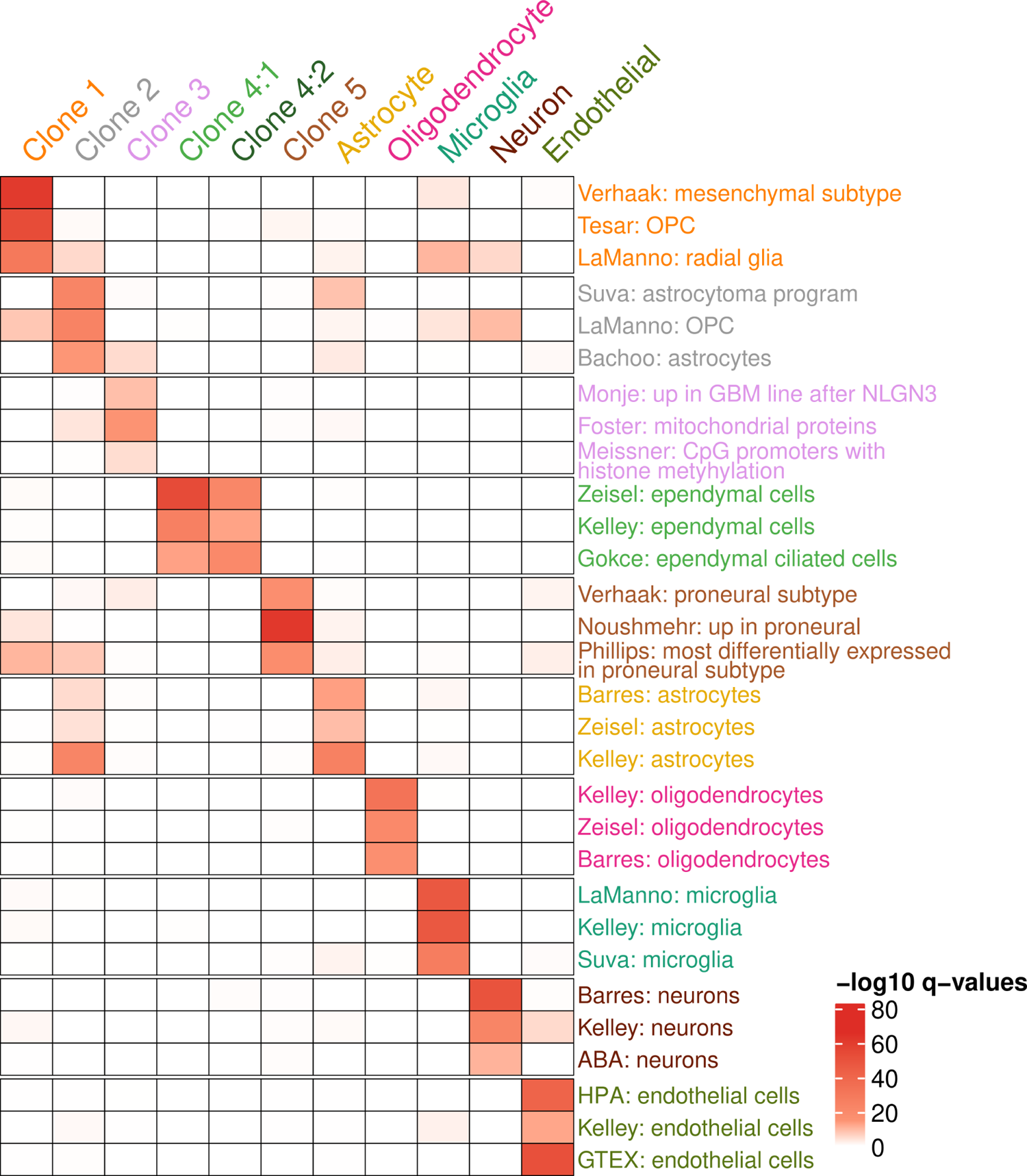
Gene set enrichment analysis supports the functional distinctness of snRNA-seq clusters. Clustered heatmap of FDR-corrected p-values (q-values) from one-sided Fisher’s exact tests comparing featured gene sets with genes that were significantly upregulated (FDR < .05) in each snRNA-seq cluster vs. all other clusters by the one-sided Wilcoxon rank-sum test.

**Figure S9.**
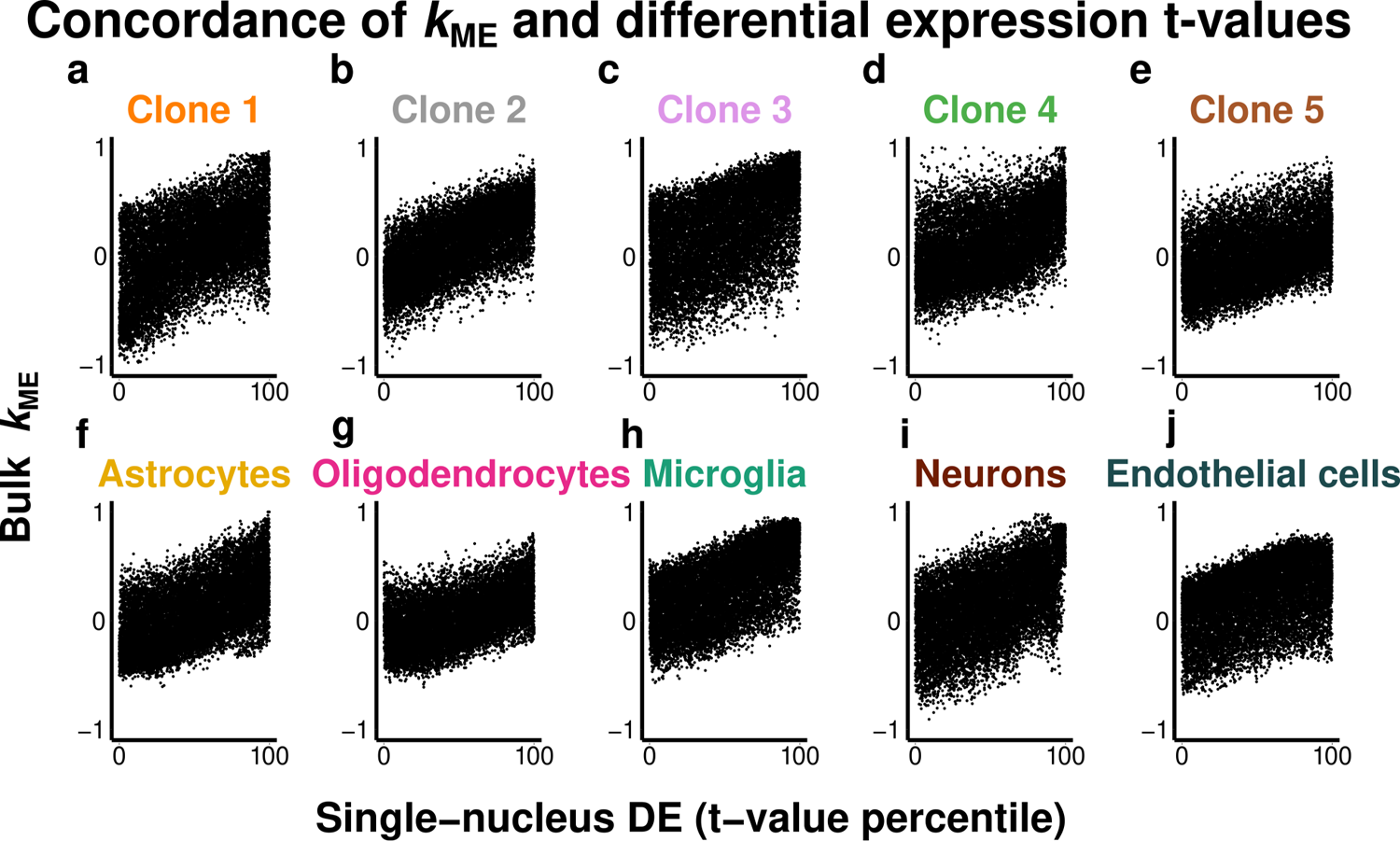
Concordance of *k*_ME_ and differential expression t-values from bulk and single-nucleus experiments. **a-j)** Differential expression (DE) t-values (calculated by t-test for all genes between each snRNA-seq cluster and all other clusters) largely predict the extent to which gene expression patterns are correlated (*k*_ME_ values) to the bulk coexpression modules most strongly associated with each clone or nonmalignant cell type.

---

**Table S1:** Section usage and quality control (case 1)

**Table S2:** Somatic mutations identified by exome sequencing (case 1)

**Table S3:** Primers for Amp-seq (case 1)

**Table S4:** Amp-seq VAFs for all tumor sections (case 1)

**Table S5:** CNV calls for all tumor sections (case 1)

**Table S6:** Cellular and clonal abundance (case 1)

**Table S7:** Module eigengenes (case 1) Table S8: *k*_ME_ values (case 1)

**Table S9:** Description of gene sets used for enrichment analysis

**Table S10:** Gene list for group-lasso model (case 1)

**Table S11:** Enrichments for group-lasso model (case 1)

**Table S12:** Section usage and quality control (normal human brain samples)

**Table S13:** Module eigengenes differential coexpression

**Table S14:** *k*_ME_ differential coexpression

**Table S15:** Section usage and quality control (case 2)

**Table S16:** Somatic mutations identified by exome sequencing (case 2)

**Table S17:** Primers for Amp-seq (case 2)

**Table S18:** Amp-seq VAFs for all tumor sections (case 2)

**Table S19:** CNV calls for all tumor sections (case 2)

**Table S20:** Cellular and clonal abundance (case 2)

**Table S21:** Single-nucleus amplicon sequencing by Mission Bio

**Table S22:** Module eigengenes (case 2)

**Table S23:** *k*_ME_ values (case 2)

**Table S24:** Gene list for lasso model (case 2)

**Table S25:** Enrichments for lasso model (case 2)

**Table S26:** Primers for snRNA-seq and snAmp-seq experiments

**Table S27:** Single-nucleus amplicon of snRNA-seq nuclei

**Table S28:** Positively differentially expressed genes for snRNA-seq

**Table S29:** Intercase correlations

## Acknowledgments

We are grateful to Brad Dispensa (UCSF), Joe Hesse (UCSF), Dirk Kleinhesselink (UCSF), Jason Jed (UCSF), and other UCSF personnel for administrative and technical support. We thank Dr. Annette Molinaro (UCSF) for statistical consultations and Dr. Anton Wellstein for helpful discussions during the project’s early stages. M.C.O. also thanks Dr. Steve Horvath, who pointed out the equivalence of correlation and the t-test when the independent variable is dichotomous. Due to space limitations, we apologize that many primary and historical publications have not been cited. This work was supported by the UCSF Program for Breakthrough Biomedical Research (M.C.O.), which is funded in part by the Sandler Foundation, a UCSF Brain Tumor SPORE Career Development Award (M.C.O.), The Shurl and Kay Curci Foundation (M.C.O.), the Dabierre Family (M.C.O.), NIH/NCI R01 CA244621 (M.C.O.), NIH/NCI T32 CA151022 (J.F.C., P.G.S.), and NIH/NINDS 1R01 NS081117 (J.J.P.).

## Author Contributions

M.C.O. conceived the approach. P.G.S. and S.J.S. reduced the approach to practice, generated all datasets, and performed most experiments and analyses. M.C.O., P.G.S., and S.J.S. wrote the manuscript. D.J.B. provided laboratory management, experimental assistance, and figure preparation. R.E. helped analyze amp-seq data. B.E.J., T.M., K.W.K, and J.F.C. analyzed WES data for case 1. D.A.L., M.B.P., and M.W.M. assisted with case selection and tumor sample acquisition. E.J.H. provided neuropathological guidance and obtained suitable control samples. M.S.B. and R.O.P. helped obtain funding and provided conceptual guidance. J.J.P. provided neuropathological guidance for both cases and performed immunohistochemistry. All authors discussed results and aided in manuscript preparation.

## Author Note

Current affiliations for co-authors include:

- SciBite, Inc., Cambridge, England (S.J.S.)
- Department of Biomedical Engineering, Oregon Health and Science University, Portland, Oregon, USA (B.H.J.)
- Dana-Farber Cancer Institute, Boston, MA, USA (T.M.)
- Department of Psychiatry, Stanford University, Palo Alto, CA, USA (K.W.K.)
- Department of Neurological Surgery, Northwestern University Feinberg School of Medicine, Chicago, USA (M.B.P.)
- Miami Neuroscience Institute, Baptist Health, Miami, FL, USA (M.W.M.)

## References

1. Burrell, R. A., McGranahan, N., Bartek, J. & Swanton, C. The causes and consequences of genetic heterogeneity in cancer evolution. Nature 501, 338–345 (2013).

2. Mazor, T., Pankov, A., Song, J. S. & Costello, J. F. Intratumoral heterogeneity of the epigenome. Cancer Cell 29, 440–451 (2016).

3. Williams, M. J., Werner, B., Barnes, C. P., Graham, T. A. & Sottoriva, A. Identification of neutral tumor evolution across cancer types. Nat. Genet. 48, 238–244 (2016).

4. Nowell, P. C. The clonal evolution of tumor cell populations. Science 194, 23–28 (1976).

5. van der Woude, L. L., Gorris, M. A. J., Halilovic, A., Figdor, C. G. & de Vries, I. J. M. Migrating into the Tumor: a Roadmap for T Cells. Trends Cancer 3, 797–808 (2017).

6. Christofides, A. et al. The complex role of tumor-infiltrating macrophages. Nat. Immunol. 23, 1148–1156 (2022).

7. Bikfalvi, A. et al. Challenges in glioblastoma research: focus on the tumor microenvironment. Trends Cancer 9, 9–27 (2023).

8. Marusyk, A., Janiszewska, M. & Polyak, K. Intratumor heterogeneity: the rosetta stone of therapy resistance. Cancer Cell 37, 471–484 (2020).

9. Vogelstein, B. et al. Cancer genome landscapes. Science 339, 1546–1558 (2013).

10. Chalmers, Z. R. et al. Analysis of 100,000 human cancer genomes reveals the landscape of tumor mutational burden. Genome Med. 9, 34 (2017).

11. Lawrence, M. S. et al. Mutational heterogeneity in cancer and the search for new cancer-associated genes. Nature 499, 214–218 (2013).

12. Martincorena, I. et al. Universal patterns of selection in cancer and somatic tissues. Cell 171, 1029–1041.e21 (2017).

13. Alexandrov, L. B. et al. The repertoire of mutational signatures in human cancer. Nature 578, 94–101 (2020).

14. Pribluda, A., de la Cruz, C. C. & Jackson, E. L. Intratumoral heterogeneity: from diversity comes resistance. Clin. Cancer Res. 21, 2916–2923 (2015).

15. Vitale, I., Shema, E., Loi, S. & Galluzzi, L. Intratumoral heterogeneity in cancer progression and response to immunotherapy. Nat. Med. 27, 212–224 (2021).

16. Bernstock, J. D. et al. Molecular and cellular intratumoral heterogeneity in primary glioblastoma: clinical and translational implications. J. Neurosurg. 1–9 (2019) doi:10.3171/2019.5.JNS19364.

17. Gerlinger, M. et al. Intratumor heterogeneity and branched evolution revealed by multiregion sequencing. N. Engl. J. Med. 366, 883–892 (2012).

18. Seol, H. et al. Intratumoral heterogeneity of HER2 gene amplification in breast cancer: its clinicopathological significance. Mod. Pathol. 25, 938–948 (2012).

19. Losi, L., Baisse, B., Bouzourene, H. & Benhattar, J. Evolution of intratumoral genetic heterogeneity during colorectal cancer progression. Carcinogenesis 26, 916–922 (2005).

20. Sottoriva, A. et al. Intratumor heterogeneity in human glioblastoma reflects cancer evolutionary dynamics. Proc Natl Acad Sci USA 110, 4009–4014 (2013).

21. Dagogo-Jack, I. & Shaw, A. T. Tumour heterogeneity and resistance to cancer therapies. Nat. Rev. Clin. Oncol. 15, 81–94 (2018).

22. Gerlovina, I., van der Laan, M. J. & Hubbard, A. Big data, small sample. Int. J. Biostat. 13, (2017).

23. Tirosh, I. et al. Single-cell RNA-seq supports a developmental hierarchy in human oligodendroglioma. Nature 539, 309–313 (2016).

24. Venteicher, A. S. et al. Decoupling genetics, lineages, and microenvironment in IDH-mutant gliomas by single-cell RNA-seq. Science 355, (2017).

25. Patel, A. P. et al. Single-cell RNA-seq highlights intratumoral heterogeneity in primary glioblastoma. Science 344, 1396–1401 (2014).

26. Young, M. D. & Behjati, S. SoupX removes ambient RNA contamination from droplet-based single-cell RNA sequencing data. Gigascience 9, giaa151. (2020).

27. Breda, J., Zavolan, M. & van Nimwegen, E. Bayesian inference of gene expression states from single-cell RNA-seq data. Nat. Biotechnol. 39, 1008–1016 (2021).

28. Zhang, M. J., Ntranos, V. & Tse, D. Determining sequencing depth in a single-cell RNA-seq experiment. Nat. Commun. 11, 774 (2020).

29. Denisenko, E. et al. Systematic assessment of tissue dissociation and storage biases in single-cell and single-nucleus RNA-seq workflows. Genome Biol. 21, 130 (2020).

30. Caglayan, E., Liu, Y. & Konopka, G. Neuronal ambient RNA contamination causes misinterpreted and masked cell types in brain single-nuclei datasets. Neuron 110, 4043–4056.e5 (2022).

31. Kelley, K. W., Nakao-Inoue, H., Molofsky, A. V. & Oldham, M. C. Variation among intact tissue samples reveals the core transcriptional features of human CNS cell classes. Nat. Neurosci. 21, 1171–1184 (2018).

32. Oldham, M. C. et al. Functional organization of the transcriptome in human brain. Nat. Neurosci. 11, 1271–1282 (2008).

33. Lui, J. H. et al. Radial glia require PDGFD-PDGFRβ signalling in human but not mouse neocortex. Nature 515, 264–268 (2014).

34. Raju, C. S. et al. Secretagogin is expressed by developing neocortical gabaergic neurons in humans but not mice and increases neurite arbor size and complexity. Cereb. Cortex 28, 1946–1958 (2018).

35. Roth, A. et al. PyClone: statistical inference of clonal population structure in cancer. Nat. Methods 11, 396–398 (2014).

36. Malikic, S., McPherson, A. W., Donmez, N. & Sahinalp, C. S. Clonality inference in multiple tumor samples using phylogeny. Bioinformatics 31, 1349–1356 (2015).

37. Louis, D. N. et al. The 2021 WHO Classification of Tumors of the Central Nervous System: a summary. Neuro Oncol. 23, 1231–1251 (2021).

38. Shen, R. & Seshan, V. E. FACETS: allele-specific copy number and clonal heterogeneity analysis tool for high-throughput DNA sequencing. Nucleic Acids Res. 44, e131 (2016).

39. Feber, A. et al. Using high-density DNA methylation arrays to profile copy number alterations. Genome Biol. 15, R30 (2014).

40. Verhaak, R. G. W. et al. Integrated genomic analysis identifies clinically relevant subtypes of glioblastoma characterized by abnormalities in PDGFRA, IDH1, EGFR, and NF1. Cancer Cell 17, 98–110 (2010).

41. Phillips, H. S. et al. Molecular subclasses of high-grade glioma predict prognosis, delineate a pattern of disease progression, and resemble stages in neurogenesis. Cancer Cell 9, 157–173 (2006).

42. Mazor, T. et al. Clonal expansion and epigenetic reprogramming following deletion or amplification of mutant IDH1. Proc Natl Acad Sci USA 114, 10743–10748 (2017).

43. Favero, F. et al. Glioblastoma adaptation traced through decline of an IDH1 clonal driver and macro-evolution of a double-minute chromosome. Ann. Oncol. 26, 880– 887 (2015).

44. Pusch, S. et al. IDH1 mutation patterns off the beaten track. Neuropathol. Appl. Neurobiol. 37, 428–430 (2011).

45. Talevich, E., Shain, A. H., Botton, T. & Bastian, B. C. CNVkit: Genome-Wide Copy Number Detection and Visualization from Targeted DNA Sequencing. PLoS Comput. Biol. 12, e1004873 (2016).

46. Eastburn, D. J., Sciambi, A. & Abate, A. R. Identification and genetic analysis of cancer cells with PCR-activated cell sorting. Nucleic Acids Res. 42, e128 (2014).

47. Rodriguez-Meira, A., O’Sullivan, J., Rahman, H. & Mead, A. J. TARGET-Seq: A Protocol for High-Sensitivity Single-Cell Mutational Analysis and Parallel RNA Sequencing. STAR Protocols 1, 100125 (2020).

48. Rodriguez-Meira, A. et al. Unravelling Intratumoral Heterogeneity through High-Sensitivity Single-Cell Mutational Analysis and Parallel RNA Sequencing. Mol. Cell 73, 1292–1305.e8 (2019).

49. Gao, R. et al. Delineating copy number and clonal substructure in human tumors from single-cell transcriptomes. Nat. Biotechnol. 39, 599–608 (2021).

50. Serin Harmanci, A., Harmanci, A. O. & Zhou, X. CaSpER identifies and visualizes CNV events by integrative analysis of single-cell or bulk RNA-sequencing data. Nat. Commun. 11, 89 (2020).

51. Street, K. et al. Slingshot: cell lineage and pseudotime inference for single-cell transcriptomics. BMC Genomics 19, 477 (2018).

52. Horvath, S. & Dong, J. Geometric interpretation of gene coexpression network analysis. PLoS Comput. Biol. 4, e1000117 (2008).

53. Szklarczyk, D. et al. STRING v11: protein-protein association networks with increased coverage, supporting functional discovery in genome-wide experimental datasets. Nucleic Acids Res. 47, D607–D613 (2019).

54. Szklarczyk, D. et al. The STRING database in 2021: customizable protein-protein networks, and functional characterization of user-uploaded gene/measurement sets. Nucleic Acids Res. 49, D605–D612 (2021).

55. Garcia-Diaz, C. et al. Glioblastoma cell fate is differentially regulated by the microenvironments of the tumor bulk and infiltrative margin. Cell Rep. 42, 112472 (2023).

56. De Silva, M. I., Stringer, B. W. & Bardy, C. Neuronal and tumourigenic boundaries of glioblastoma plasticity. Trends Cancer 9, 223–236 (2023).

57. Dirkse, A. et al. Stem cell-associated heterogeneity in Glioblastoma results from intrinsic tumor plasticity shaped by the microenvironment. Nat. Commun. 10, 1787 (2019).

58. Spassky, N. et al. Adult ependymal cells are postmitotic and are derived from radial glial cells during embryogenesis. J. Neurosci. 25, 10–18 (2005).

59. Redmond, S. A. et al. Development of Ependymal and Postnatal Neural Stem Cells and Their Origin from a Common Embryonic Progenitor. Cell Rep. 27, 429–441.e3 (2019).

60. Hahn, W. C. et al. An expanded universe of cancer targets. Cell 184, 1142–1155 (2021).

61. Penning, T. M. AKR1C3 (type 5 17β-hydroxysteroid dehydrogenase/prostaglandin F synthase): Roles in malignancy and endocrine disorders. Mol. Cell. Endocrinol. 489, 82–91 (2019).

62. Zhou, Q. et al. A Positive Feedback Loop of AKR1C3-Mediated Activation of NF-κB and STAT3 Facilitates Proliferation and Metastasis in Hepatocellular Carcinoma. Cancer Res. 81, 1361–1374 (2021).

63. Liu, C. et al. Intracrine androgens and AKR1C3 activation confer resistance to enzalutamide in prostate cancer. Cancer Res. 75, 1413–1422 (2015).

64. Bortolozzi, R. et al. AKR1C enzymes sustain therapy resistance in paediatric T-ALL. Br. J. Cancer 118, 985–994 (2018).

65. Dang, L. et al. Cancer-associated IDH1 mutations produce 2-hydroxyglutarate. Nature 462, 739–744 (2009).

66. Xu, W. et al. Oncometabolite 2-hydroxyglutarate is a competitive inhibitor of α-ketoglutarate-dependent dioxygenases. Cancer Cell 19, 17–30 (2011).

67. Mellinghoff, I. K. et al. Vorasidenib in IDH1- or IDH2-Mutant Low-Grade Glioma. N. Engl. J. Med. 389, 589–601 (2023).

68. Faul, F., Erdfelder, E., Buchner, A. & Lang, A.-G. Statistical power analyses using G*Power 3.1: Tests for correlation and regression analyses. Behav. Res. Methods 41, 1149–1160 (2009).

69. Langfelder, P. & Horvath, S. WGCNA: an R package for weighted correlation network analysis. BMC Bioinformatics 9, 559 (2008).

70. Langfelder, P. & Horvath, S. Fast R Functions for Robust Correlations and Hierarchical Clustering. J. Stat. Softw. 46, (2012).

71. Johnson, B. E. et al. Mutational analysis reveals the origin and therapy-driven evolution of recurrent glioma. Science 343, 189–193 (2014).

72. Li, H. & Durbin, R. Fast and accurate long-read alignment with Burrows-Wheeler transform. Bioinformatics 26, 589–595 (2010).

73. Broad Institute, G. Repository. Picard Toolkit. (Broad Institute, 2019).

74. Van der Auwera, G. A. et al. From FastQ data to high confidence variant calls: the Genome Analysis Toolkit best practices pipeline. Curr. Protoc. Bioinformatics 11, 11.10.1–11.10.33 (2013).

75. Cibulskis, K. et al. Sensitive detection of somatic point mutations in impure and heterogeneous cancer samples. Nat. Biotechnol. 31, 213–219 (2013).

76. Ye, K., Schulz, M. H., Long, Q., Apweiler, R. & Ning, Z. Pindel: a pattern growth approach to detect break points of large deletions and medium sized insertions from paired-end short reads. Bioinformatics 25, 2865–2871 (2009).

77. Wang, K., Li, M. & Hakonarson, H. ANNOVAR: functional annotation of genetic variants from high-throughput sequencing data. Nucleic Acids Res. 38, e164 (2010).

78. Sherry, S. T. et al. dbSNP: the NCBI database of genetic variation. Nucleic Acids Res. 29, 308–311 (2001).

79. 1000 Genomes Project Consortium et al. A global reference for human genetic variation. Nature 526, 68–74 (2015).

80. McLaren, W. et al. The ensembl variant effect predictor. Genome Biol. 17, 122 (2016).

81. Ye, J. et al. Primer-BLAST: a tool to design target-specific primers for polymerase chain reaction. BMC Bioinformatics 13, 134 (2012).

82. Li, H. Aligning sequence reads, clone sequences and assembly contigs with BWA-MEM. arXiv (2013) doi:10.48550/arxiv.1303.3997.

83. Li, H. et al. The Sequence Alignment/Map format and SAMtools. Bioinformatics 25, 2078–2079 (2009).

84. Koboldt, D. C. et al. VarScan 2: somatic mutation and copy number alteration discovery in cancer by exome sequencing. Genome Res. 22, 568–576 (2012).

85. Thorndike, R. L. Who belongs in the family? Psychometrika 18, 267–276 (1953).

86. Rousseeuw, P. J. Silhouettes: A graphical aid to the interpretation and validation of cluster analysis. Journal of Computational and Applied Mathematics 20, 53–65 (1987).

87. Maechler, M., Rousseeuw, P., Struyf, A., Hubert, M. & Hornik, K. cluster: Cluster Analysis Basics and Extensions. Available online at: https://CRAN.R-project.org/package=cluster. (2022).

88. Morris, T. J. et al. Champ: 450k chip analysis methylation pipeline. Bioinformatics 30, 428–430 (2014).

89. Aryee, M. J. et al. Minfi: a flexible and comprehensive Bioconductor package for the analysis of Infinium DNA methylation microarrays. Bioinformatics 30, 1363–1369 (2014).

90. Fortin, J.-P., Triche, T. J. & Hansen, K. D. Preprocessing, normalization and integration of the Illumina HumanMethylationEPIC array with minfi. Bioinformatics 33, 558–560 (2017).

91. Zhou, W., Laird, P. W. & Shen, H. Comprehensive characterization, annotation and innovative use of Infinium DNA methylation BeadChip probes. Nucleic Acids Res. 45, e22 (2017).

92. Dedeurwaerder, S. et al. Evaluation of the Infinium Methylation 450K technology. Epigenomics 3, 771–784 (2011).

93. Teschendorff, A. E. et al. A beta-mixture quantile normalization method for correcting probe design bias in Illumina Infinium 450 k DNA methylation data. Bioinformatics 29, 189–196 (2013).

94. Oldham, M. C., Langfelder, P. & Horvath, S. Network methods for describing sample relationships in genomic datasets: application to Huntington’s disease. BMC Syst. Biol. 6, 63 (2012).

95. Bolstad, B. M., Irizarry, R. A., Astrand, M. & Speed, T. P. A comparison of normalization methods for high density oligonucleotide array data based on variance and bias. Bioinformatics 19, 185–193 (2003).

96. Johnson, W. E., Li, C. & Rabinovic, A. Adjusting batch effects in microarray expression data using empirical Bayes methods. Biostatistics 8, 118–127 (2007).

97. Barbosa-Morais, N. L. et al. A re-annotation pipeline for Illumina BeadArrays: improving the interpretation of gene expression data. Nucleic Acids Res. 38, e17 (2010).

98. Andrews, S., et al. FastQC: a quality control tool for high throughput sequence data. Available online at: http://www.bioinformatics.babraham.ac.uk/projects/fastqc. (2010).

99. Martin, M. Cutadapt removes adapter sequences from high-throughput sequencing reads. EMBnet j. 17, 10 (2011).

100. Langmead, B. & Salzberg, S. L. Fast gapped-read alignment with Bowtie 2. Nat. Methods 9, 357–359 (2012).

101. Lander, E. S. et al. Initial sequencing and analysis of the human genome. Nature 409, 860–921 (2001).

102. Rosenbloom, K. R. et al. ENCODE whole-genome data in the UCSC Genome Browser: update 2012. Nucleic Acids Res. 40, D912–7 (2012).

103. Risso, D., Ngai, J., Speed, T. P. & Dudoit, S. Normalization of RNA-seq data using factor analysis of control genes or samples. Nat. Biotechnol. 32, 896–902 (2014).

104. Olshen, A. B. et al. Parent-specific copy number in paired tumor-normal studies using circular binary segmentation. Bioinformatics 27, 2038–2046 (2011).

105. Venkatraman, E. S. & Olshen, A. B. A faster circular binary segmentation algorithm for the analysis of array CGH data. Bioinformatics 23, 657–663 (2007).

106. Glur, C. data.tree: General Purpose Hierarchical Data Structure. Available online at: https://CRAN.R-project.org/package=data.tree. (2020).

107. Iannone, R. DiagrammeR: Graph/Network Visualization. Available online at: https://CRAN.R-project.org/package=DiagrammeR. (2022).

108. Storey, J. D. & Tibshirani, R. Statistical significance for genomewide studies. Proc Natl Acad Sci USA 100, 9440–9445 (2003).

109. Klosa, J., Simon, N., Westermark, P. O., Liebscher, V. & Wittenburg, D. Seagull: lasso, group lasso and sparse-group lasso regularization for linear regression models via proximal gradient descent. BMC Bioinformatics 21, 407 (2020).

110. Tibshirani, R. Regression Shrinkage and Selection via the Lasso. J R Stat Soc Series B Stat Methodol 58, 267–288 (1995).

111. Yuan, M. & Lin, Y. Model selection and estimation in regression with grouped variables. J. Royal Statistical Soc. B 68, 49–67 (2006).

112. Laurin, C., Boomsma, D. & Lubke, G. The use of vector bootstrapping to improve variable selection precision in Lasso models. Stat. Appl. Genet. Mol. Biol. 15, 305–320 (2016).

113. Mason, M. J., Fan, G., Plath, K., Zhou, Q. & Horvath, S. Signed weighted gene co-expression network analysis of transcriptional regulation in murine embryonic stem cells. BMC Genomics 10, 327 (2009).

114. Tesson, B. M., Breitling, R. & Jansen, R. C. DiffCoEx: a simple and sensitive method to find differentially coexpressed gene modules. BMC Bioinformatics 11, 497 (2010).

115. Smith, T., Heger, A. & Sudbery, I. UMI-tools: modeling sequencing errors in Unique Molecular Identifiers to improve quantification accuracy. Genome Res. 27, 491– 499 (2017).

116. Bolger, A. M., Lohse, M. & Usadel, B. Trimmomatic: A flexible trimmer for Illumina sequence data. Bioinformatics 30, 2114–2120 (2014).

117. Dobin, A. et al. STAR: ultrafast universal RNA-seq aligner. Bioinformatics 29, 15– 21 (2013).

118. Liao, Y., Smyth, G. K. & Shi, W. featureCounts: an efficient general purpose program for assigning sequence reads to genomic features. Bioinformatics 30, 923–930 (2014).

119. Harrow, J. et al. GENCODE: the reference human genome annotation for The ENCODE Project. Genome Res. 22, 1760–1774 (2012).

120. Melville, J. uwot: The Uniform Manifold Approximation and Projection (UMAP) Method for Dimensionality Reduction. (2021).

121. Untergasser, A. et al. Primer3--new capabilities and interfaces. Nucleic Acids Res. 40, e115 (2012).

122. Frank, D. N. BARCRAWL and BARTAB: software tools for the design and implementation of barcoded primers for highly multiplexed DNA sequencing. BMC Bioinformatics 10, 362 (2009).

123. Butts, C. T. network: A Package for Managing Relational Data inR. J. Stat. Softw. 24, (2008).

124. Bojanowski, M. intergraph: Coercion Routines for Network Data Objects. Available online at: http://mbojan.github.io/intergraph. (2015).

125. Briatte, F. ggnetwork: Geometries to Plot Networks with “ggplot2”. Available online at: https://CRAN.R-project.org/package=ggnetwork. (2021).

126. Karlsson, M. et al. A single-cell type transcriptomics map of human tissues. Sci. Adv. 7, (2021).

127. Wickham, H. ggplot2: Elegant Graphics for Data Analysis (Use R!). 276 (Springer, 2016).

128. Dowle, M. & Srinivasan, A. data.table: Extension of ‘data.framè. Available online at: https://CRAN.R-project.org/package=data.table. (2021).

129. Neuwirth, E. RColorBrewer: ColorBrewer Palettes. Available online at: https://CRAN.R-project.org/package=RColorBrewer. (2022).

130. Auguie, B. gridExtra: Miscellaneous Functions for “Grid” Graphics. Available online at: https://CRAN.R-project.org/package=gridExtra. (2017).

131. Gu, Z., Eils, R. & Schlesner, M. Complex heatmaps reveal patterns and correlations in multidimensional genomic data. Bioinformatics 32, 2847–2849 (2016).

132. Gu, Z., Gu, L., Eils, R., Schlesner, M. & Brors, B. circlize Implements and enhances circular visualization in R. Bioinformatics 30, 2811–2812 (2014).

133. Ahlmann-Eltze, C. & Patil, I. ggsignif: R Package for Displaying Significance Brackets for “ggplot2.” (2021) doi:10.31234/osf.io/7awm6.

